# Mechanistic Basis for the Selective Recognition of the Fcγ Receptor IIa by Monoclonal Antibody IV.3

**DOI:** 10.64898/2026.03.05.709909

**Authors:** Jialing Wang, Sabina Novack, Jihong Li, Emily G. Niejadlik, Stylianos Bournazos, Barry S. Coller, Marta Filizola

## Abstract

The monoclonal antibody IV.3 selectively binds the platelet Fcγ receptor IIa (FcγRIIa), potently blocking immune complex engagement without cross–reacting with the closely–related FcγRIIb. This specificity has made IV.3 invaluable for dissecting FcγRIIa–mediated activation in diverse conditions, including infection, autoimmunity, thromboinflammation, and platelet–mediated thrombosis. We combined cryogenic electron microscopy, surface plasmon resonance, alchemical free energy calculations, and molecular dynamics simulations to elucidate IV.3’s binding sites on FcγRIIa and the mechanistic basis of IV.3 specificity. We find that IV.3 engages a broader FcγRIIa epitope than previously recognized, extending beyond residues H/R134 and L135 (R and S in FcγRIIb). Simulations of FcγIIa–R134 variants bearing either L135 or S135 reveal that IV.3 specificity arises from hydrophobic stabilization mediated by L135 and disruption of an R134-specific interaction network in the presence of S135. These findings provide a mechanistic framework for rational design of FcγRIIa-targeted therapeutics.

## Introduction

Fc gamma receptor IIa (FcγRIIa) is a cell–surface receptor that recognizes the fragment crystallizable (Fc) region of immunoglobulin G (IgG) antibodies and transmits this interaction across the cell membrane, initiating intracellular signaling via its cytoplasmic immunoreceptor tyrosine–based activation motif (ITAM).^1^ Although FcγRIIa has low affinity for monomeric IgG, it binds immune complexes with much higher affinity, promoting receptor clustering and downstream signaling that is important in platelet activation related to infection and autoimmunity, in some cases leading to thromboinflammation and pathological thrombosis.^1^ Notably, while platelet FcγRIIa has been implicated in enhancing platelet responses to activation, it is not required for hemostasis.^1^

The murine IgG2b monoclonal antibody (mAb) IV.3, developed nearly four decades ago,^2^ potently blocks IgG Fc engagement to FcγRIIa and has played a vital role in defining the biological and pathological roles of this receptor in both in vitro and in vivo experiments,^3, 4, 5, 6^ with PubMed currently listing more than 150 publications documenting its use in a wide variety of studies. A humanized version of mAb IV.3 is currently under development as a therapeutic.^7^

A common FcγRIIa polymorphism (H/R; rs1801274), originally mapped to residue 131 and later reassigned to residue 134,^8, 9^ alters IgG Fc affinity. The mAb IV.3 recognizes both polymorphic variants,^10^ further increasing its utility as a reagent for receptor identification, quantification, and blockade. Notably, IV.3 has been reported to be highly selective for FcγRIIa and does not cross–react with the closely–related receptor FcγRIIb, which is expressed on neutrophils, eosinophils, monocytes, macrophages, dendritic cells, and B cells, but not on platelets.^9, 11, 12^

Despite its extensive use, the molecular basis for the specificity of IV.3 has remained incompletely resolved. Early chimeric receptor and site–directed mutagenesis experiments localized the FcγRIIa functional epitope for IV.3 to a short segment (residues 132–137) that also encompasses the common FcγRIIa polymorphism at position 134 (H/R). These experiments identified residue 135 as a key determinant distinguishing FcγRIIa from FcγRIIb:^9^ substituting FcγRIIa L135 with the corresponding serine residue of FcγRIIb abolished IV.3 binding to CHO cells expressing FcγRIIa, whereas substituting neighboring FcγRIIb–specific residues at positions 130 and 138 had no effect.^9^ Yet, epitope mapping by domain swaps could not capture the full three–dimensional architecture of the antibody–receptor interface or the role of cooperative loop networks that often shape antibody specificity. Moreover, while IV.3 is known to recognize both H134 and R134 variants,^10^ the structural and energetic basis of their interactions, and how those interactions might affect local conformational dynamics at the interface, have not been defined at atomic resolution.

A high–resolution structural and mechanistic description is needed for two reasons. First, FcγRIIa is a validated driver of immune complex–mediated platelet activation, making it an attractive therapeutic target in thromboinflammatory disorders where selective antagonism of FcγRIIa without perturbing FcγRIIb’s important immunoregulatory functions is desirable.^13^ Second, IV.3 itself, and derivatives engineered to minimize Fc effector activity, have been explored preclinically and clinically as therapeutic inhibitors of FcγRIIa.^7^ Rational improvement of such agents requires a detailed blueprint of the interactions that confer potency and selectivity, as well as an understanding of how common polymorphisms modulate these interactions through both static contacts and conformational dynamics.

Here we address these knowledge gaps by combining cryogenic electron microscopy (cryo–EM), surface plasmon resonance (SPR), alchemical free energy calculations (FEC), and molecular dynamics (MD) simulations. We determine a 3.5 Å structure of the FcγRIIa–IV.3 Fab complex (FcγRIIa–H134 variant), which reveals an extended binding interface involving three receptor loops, defines the contributions of residues H134 and L135 alongside additional interaction hubs such as P117, V119, and Y160, and shows how IV.3 sterically occludes the IgG–binding surface. SPR measurements confirm that IV.3 binds both FcγRIIa polymorphic variants with subnanomolar affinity, while exhibiting negligible binding to FcγRIIb. Integrated alchemical free energy and MD analyses establish that L135 provides critical hydrophobic stabilization within the FcγRIIa–IV.3 interface, while the FcγRIIa–R134 variant introduces distinct interactions facilitated by a rearrangement of the IV.3 loop spanning residues D119–D123, with the dominant contribution arising from a Y122–Y51 gating–like mechanism. In contrast, substitution of L135 with serine disrupts this interaction network, and the FcγRIIb–mimicking R134/S135 construct appears to synergistically destabilize binding. Together, these findings define the structural and dynamic determinants underlying IV.3 specificity for FcγRIIa and provide a mechanistic foundation for the rational design of selective FcγRIIa–targeted therapeutics.

## Results

### Overall structure of the FcγRIIa–IV.3 Fab complex

To structurally characterize the interaction between the FcγRIIa–H134 polymorphic variant and mAb IV.3, an aliquot of the FcγRIIa–IV.3 Fab complex prepared for negative–stain EM analysis (see Methods and Supplementary Fig. 1) was vitrified and cryo–EM data were collected on a Titan Krios electron microscope equipped with a K3 direct electron detector. Image processing yielded a reconstruction of the FcγRIIa–IV.3 Fab complex at an overall resolution of 3.5 Å, with up to 3.0 Å local resolution at the interface between IV.3 and FcγRIIa (Fig. 1, Supplementary Figs. 2 and 3; Supplementary Table 1).

**Fig. 1.**
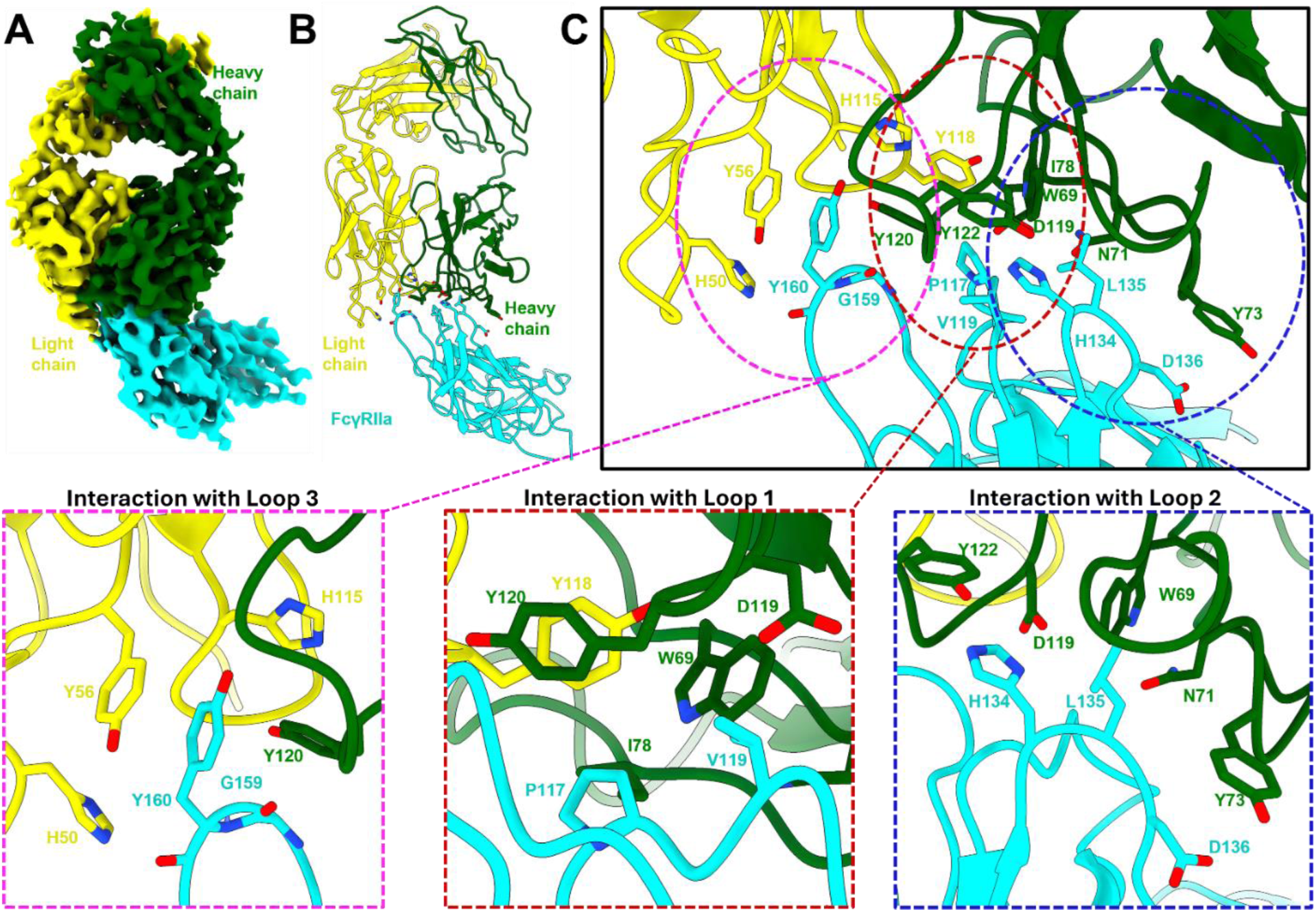
The cryo–EM structure of the FcγRIIa–H134 variant in complex with IV.3 Fab. **(A)** Density map of the FcγRIIa–IV.3 Fab complex at 3.5–Å resolution, colored by subunit and labeled accordingly. **(B)** Cartoon representation of the atomic model of the FcγRIIa–IV.3 Fab complex, colored by subunit and labeled accordingly. **(C) Top:** Enlarged view of the receptor–antibody interface, with FcγRIIa shown in cyan and the IV.3 Fab heavy and light chains in green and yellow, respectively. Residues identified as interacting by ProLIF interaction–fingerprint analysis (see Supplementary Table 2 for interaction details) are shown as sticks and labeled. **Bottom:** Close–up views of key interaction hubs within the three FcγRIIa loops — from left to right, residues 156–163 (loop 3), 113–120 (loop 1), and 133–137 (loop 2) — shown as sticks and labeled.

The cryo–EM structure revealed that the IV.3 Fab engages FcγRIIa (A chain) through both its heavy (H) and light (L) chains (Fig. 1). The refined atomic model, including residues at the interface, fits well into the cryo–EM density map (Supplementary Fig. 3), allowing for an unambiguous assignment of most interacting residues. The structure identifies three FcγRIIa loop regions at the binding interface, residues 113–120 (loop1), 133–137 (loop2), and 156–163 (loop3), defined by the presence of residues located within 4–5 Å of the mAb IV.3 Fab (Fig. 1, Supplementary Fig. 4).

Protein–protein interaction analysis of the FcγRIIa–H134–IV.3 cryo–EM structure with ProLIF^14^ (Supplementary Table 2) reveals that residues within loop 1 form extensive hydrophobic interactions with the IV.3 Fab. In particular, P117 forms hydrophobic contacts with IV.3 Fab heavy–chain residues W69, I78, and Y120, as well as with light–chain residue Y118, and additionally engages in van der Waals (VdW) interaction with IV.3 heavy–chain residue I78 and light–chain residue Y118 (Fig. 1). FcγRIIa loop 1 residue V119 further contributes to the binding interface through a hydrophobic interaction with IV.3 heavy–chain residue Y120 and VdW interaction with IV.3 heavy–chain residue D119. Protein–protein interaction analysis of our deposited FcγRIIa–H134–IV.3 cryo–EM structure indicates that no additional loop 1 residues directly engage IV.3. While the density for K120 is compatible with an alternative side-chain orientation toward IV.3 residue G121 that could permit a hydrogen bond with the G121 backbone carbonyl, our MD simulations (see below) indicate that this interaction is transient and therefore unlikely to be functionally relevant. Loop 2, which encompasses the core epitope region (residues 132–137), interacts predominantly with the IV.3 Fab heavy chain. FcγRIIa H134 forms both hydrophobic and VdW interactions with IV.3 heavy–chain residue Y122, along with an additional VdW contact with IV.3 heavy–chain residue D119. Within this FcγRIIa loop, L135 engages in a hydrophobic interaction with IV.3 heavy chain W69 and establishes additional VdW interactions with IV.3 heavy–chain residues N71 and Y73, while FcγRIIa D136 forms a hydrophobic interaction with IV.3 heavy chain Y73 (Fig. 1). FcγRIIa loop 3 appears to contribute a major stabilizing interaction centered on Y160. This residue forms hydrophobic interactions with IV.3 Fab heavy chain Y120, as well as light–chain residues H50, Y56, and H115. In addition, FcγRIIa Y160 forms a hydrogen bond through its backbone carbonyl oxygen with the Nε2 atom of the IV.3 light chain H50 imidazole ring, accompanied by VdW interactions with both IV.3 light–chain residues H50 and Y56. Notably, the side chain aromatic ring of IV.3 light chain Y56 also participates in face–to–face and π–π stacking interactions with FcγRIIa Y160 (Fig. 1; Supplementary Table 2).

The reported crystal structure of the FcγRIIa–R134 variant in complex with the Fc domain of human IgG_1_ (PDB code ID: 3RY6^9^) reveals an interaction interface that substantially overlaps with that observed for mAb IV.3, providing a structural explanation for IV.3’s ability to block IgG binding to the receptor (Supplementary Table 2 and Supplementary Fig. 5C). Several FcγRIIa residues, including Y160, P117, V119, and the polymorphic residue R134, contribute directly to binding in both complexes (compare insets in Supplementary Figs. 5A and 5B). In contrast, FcγRIIa residues W90, W113, K120, and F132 are uniquely involved in the interaction of FcγRIIa–R134 with the IgG1 Fc domain, whereas FcγRIIa L135, D136, and G159 are specific to the interaction of the FcγRIIa–H134 variant with mAb IV.3 (Supplementary Table 2 and Supplementary Figs. 5A and 5B).

### Thermodynamics of mAb IV.3 binding specificity for FcγRIIa

To define the thermodynamic basis of mAb VI.3’s binding specificity for FcγRIIa, we constructed coupled thermodynamic cycles (Supplementary Fig. 6B–6D) that provide the theoretical framework for alchemical free–energy calculations quantifying the individual and cooperative effects of the H134R and L135S transformations, introduced singly or in combination to mimic the FcγRIIb epitope, on the binding energetics of the FcγRIIa–IV.3 Fab complex resolved by cryo–EM.

State–wise relative free–energy differences (ΔG) arising from amino acid substitution in FcγRIIa, both in solvent and in complex with the mAb IV.3 Fab, were computed using the expanded ensemble (EE) method.^15^ Supplementary Fig. 6E shows the resulting free energy trajectories and Wang–Landau increment trajectories. The calculated ΔΔG for the double H134R/L135S alchemical transformation in FcγRIIa is 95.34 ± 0.01 kcal mol⁻¹, indicating a pronounced destabilization of IV.3 binding to this FcγRIIa construct, which mimics the FcγRIIb sequence in the region of IV.3 binding (Supplementary Table 3). This result is consistent with the profound difference in mAb IV.3 binding to FcγRIIa–R134 versus FcγRIIb that we measured by SPR (Supplementary Fig. 7), showing subnanomolar affinity for FcγRIIa–R134 (K_D_ = 4.76 x 10^-10^ M) and no detectable binding to FcγRIIb (K_D_ >10^-6^ M), in agreement with previously reported binding data.^9^ In contrast, the calculated ΔΔG value for the H134R transformation within FcγRIIa was –0.84 ± 0.05 kcal mol⁻¹, consistent with the minimal enhancement in FcγRIIa–IV.3 Fab binding affinity observed by SPR (Supplementary Fig. 7), where FcγRIIa–R134 (K_D_ = 4.76 x 10^-10^ M) binds only slightly more tightly than FcγRIIa–H134 (K_D_ = 7.87 x 10^-10^ M). The ΔΔG value for the L135S substitution (0.59 ± 0.01 kcal mol⁻¹; Supplementary Table 3) indicates a slight destabilization of IV.3 Fab binding, consistent with prior binding data for this mutant receptor.^9^ To assess potential sampling bottlenecks in interfacial conformational dynamics that may arise from initiating alchemical transformations from the cryo–EM structure of the FcγRIIa–H134 variant, and to more effectively probe the mechanistic basis of FcγRIIa recognition by IV.3, we next performed MD simulations.

### Interaction dynamics driving FcγRIIa–IV.3 specificity

We carried out standard MD simulations of the cryo–EM structure of the FcγRIIa–H134 variant in complex with IV.3 Fab, along with simulations of structurally analogous models in which H134 was substituted with an arginine (R134 variant) or in which both H134 and L135 were substituted with arginine and serine, respectively (R134/S135 system), to mimic the FcγRIIb interface. To compare the conformational dynamics of the FcγRIIa–IV.3 interface across these three systems, we conducted time–lagged independent component analysis (tICA),^16, 17^ using a shared feature set. This set included Cα_i_–Cα_j_ and Cβ_i_–Cβ_j_ distances and Cα_i_–Cβ_i_–Cβ_j_–Cα_j_ torsion angles for residue pairs forming heavy–atom contacts (≤4.5 Å) with interfacial positions Y160, 134, and 135, as well as K120. K120 was included based on its involvement in an anionic interaction in the crystal structure of the FcγRIIa–R134 variant bound to the Fc domain of human IgG_1_ (PDB code ID: 3RY6^9^) as well as the ambiguity in its side-chain orientation inferred from the cryo-EM density of the FcγRIIa–H134–IV.3 Fab complex (Supplementary Table 2). The shared feature set also incorporated the backbone root mean square deviation (RMSD) relative to the FcγRIIa–IV.3 Fab cryo–EM structure. As shown in Fig. 2A, each system occupies a distinct region of tICA space, indicating that the H134, R134, and R134/S135 variants of the FcγRIIa–IV.3 Fab complex explore unique conformational ensembles at the FcγRIIa–IV.3 interface.

**Fig. 2.**
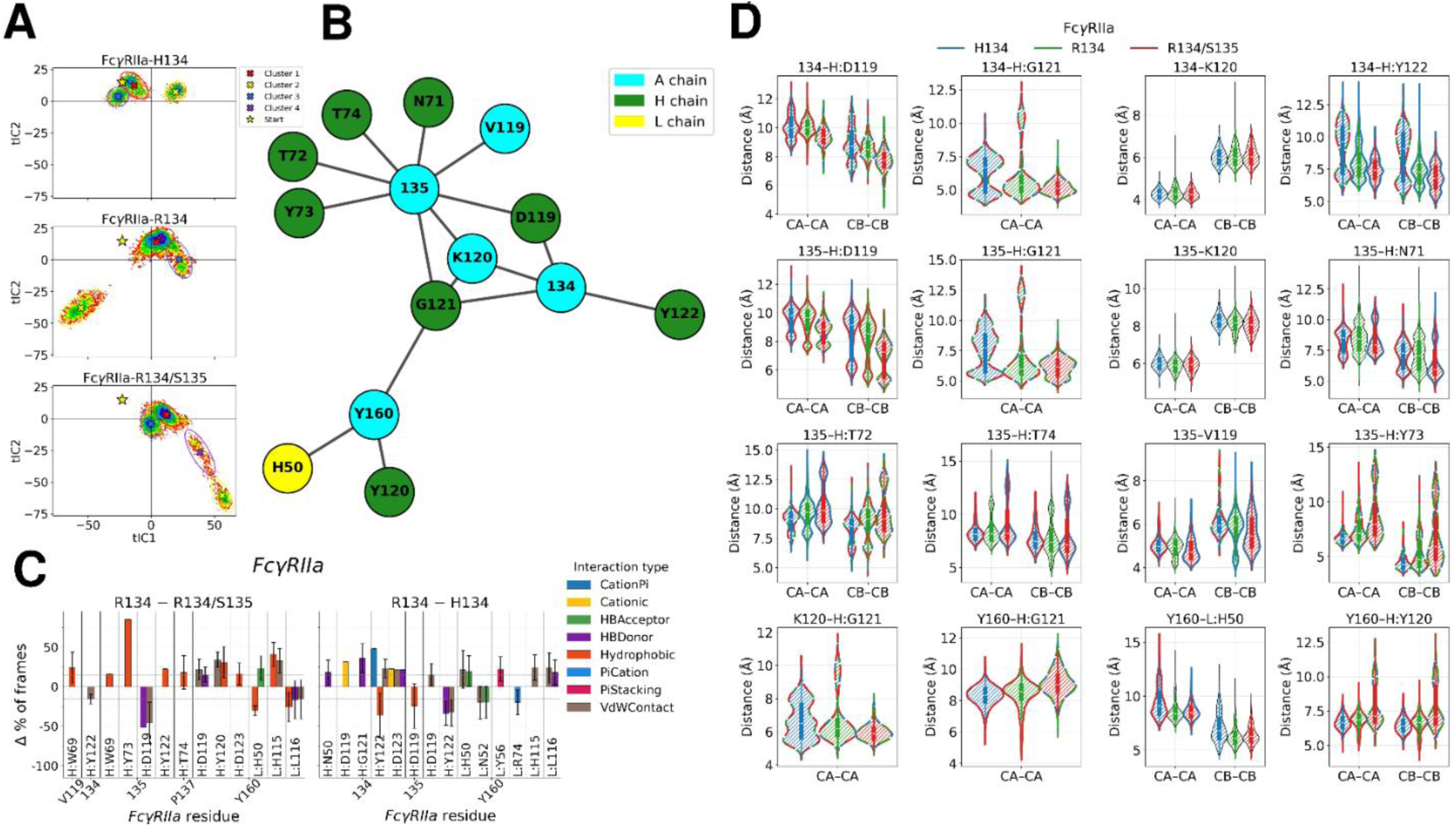
Analysis of the FcγRIIa–IV.3 Fab interface dynamics. **(A)** Time–lagged independent component analysis (tICA) projections of the dominant tICs for the FcγRIIa H134, R134, and R134/S135 constructs. The yellow star denotes the conformation of the FcƔRIIa–IV.3 Fab cryo–EM structure. Projections of the dominant tICs for the FcγRIIa H134, R134, and R134/S135 constructs were clustered into 3, 3, and 4 states, respectively, denoted by colored ovals (cluster 1: red, cluster 2: yellow, cluster 3: blue, cluster 4: purple), with the corresponding colored “X” indicating each cluster center. Horizontal and vertical black lines mark x=0 and y=0; **(B)** Time–lagged covariance network highlighting the most highly connected, heavily weighted feature “path,” as identified by a greedy algorithm and coarse–grained to chain:residue resolution. Nodes are colored by chain [cyan: FcγRIIa (A chain); green: heavy chain (H chain); yellow: light chain (L chain)] and residue index; **(C)** ProLIF–derived protein–protein interaction fingerprints showing differences in interaction frequency (>15%) between the simulated FcγRIIa R134 and R134/S135 systems (left) and between the FcγRIIa R134 and H134 systems (right). Interacting IV.3 residues are annotated as H:residue number or L:residue number, indicating residues from the heavy and light chains, respectively; **(D)** Violin plots of the feature distributions along the dominant covariance–network path. Significant differences between distributions were determined using the Jensen–Shannon divergence (threshold > 0.07 nats). The outline color indicates the construct with which the feature distribution significantly differs from (blue: H134; green: R134; red: R134/S135). Dashed two–color outlines indicate significant differences with both other constructs, whereas black outlines indicate no significant differences. Interacting FcγRIIa residues are labeled by residue number, whereas IV.3 residues are annotated as H:residue number or L:residue number, indicating residues from the heavy and light chains, respectively.

Using the time–lagged covariance matrix, we constructed a time–lagged covariance network of dynamically coupled features (Fig. 2B) and applied a greedy most–heavily–weighted path algorithm^18^ to identify a subnetwork of residues at the FcγRIIa–IV.3 interface exhibiting the strongest covariation. This analysis revealed that FcγRIIa residues at position 134 covary primarily with the adjacent FcγRIIa residue K120 and IV.3 heavy–chain residues Y122, D119, and G121, with FcγRIIa K120 serving as the principal connector between residues at positions 134 and 135. In contrast, FcγRIIa residues at position 135 participate in a broader local covariation network involving adjacent FcγRIIa loop residues V119 and K120 together with IV.3 heavy–chain residues D119, G121, Y73, T72, T74, and N71 (Fig. 2B). FcγRIIa residue Y160 was the only position showing coordinated motion with residues on both IV.3 light (H50) and heavy (Y120, G121) chains (Fig. 2B).

Analysis of protein–protein interaction differences in MD simulations of the R134, H134, and R134/S135 variants of the FcγRIIa–IV.3 Fab complex (Fig. 2C) provided further mechanistic insight into epitope discrimination. Comparison between simulations of the R134 and R134/S135 variants (Fig. 2C; left plot) revealed that, on the R134 background, L135 forms hydrophobic interactions with IV.3 heavy–chain residues W69, Y73, and Y122, occurring most frequently (85%) with Y73. These interactions are absent in simulations of the R134/S135 variant, which mimics FcγRIIb. Instead, in this variant, S135 forms hydrogen bonds and van der Waals interactions with IV.3 heavy–chain residue D119 (Fig. 2C; left plot), potentially interfering with R134’s ability to engage in electrostatic interactions with this residue. As a result, R134 reorients to form van der Waals contacts with IV.3 heavy–chain residue Y122 (Fig. 2C, left plot), which diminishes the strength of the epitope–specific interactions.

FcγRIIa residue Y160 functions as an interaction hub across all simulated systems. In the R134 variant with leucine at position 135, Y160 forms its most persistent contacts (>25%) with IV.3 heavy–chain residue Y120 and light–chain residue H115 (Fig. 2C; left plot), whereas in the R134/S135 variant, these interactions shift to light–chain residues H50 and L116 (Fig. 2C; left plot). Comparison between simulations of the R134 and H134 variants further reveals a direct effect of the R134 substitution on the Y160 interaction fingerprint: hydrogen–bonding and vdW interactions to IV.3 heavy–chain Y122 are lost, while new vdW interactions with IV.3 heavy–chain D119 and IV.3 light–chain H115, H50, and L116 emerge, together with polar interactions involving H50 and L116, and π–π stacking involving Y56 (Fig. 2C; right plot). Notably, simulations of the R134 variant of the FcγRIIa–IV.3 Fab complex showed that formation of this R134–specific interaction network is enabled by a rearrangement of the IV.3 D119–D123 loop, which promotes a IV.3 Y122–Y51 gating–like motion: IV.3 Y122 flips over to encapsulate FcγRIIa R134, thereby establishing the aforementioned specific interactions (see description of Figs. 3B, 3C, below, as well as in Supplementary Movie 1).

**Fig. 3.**
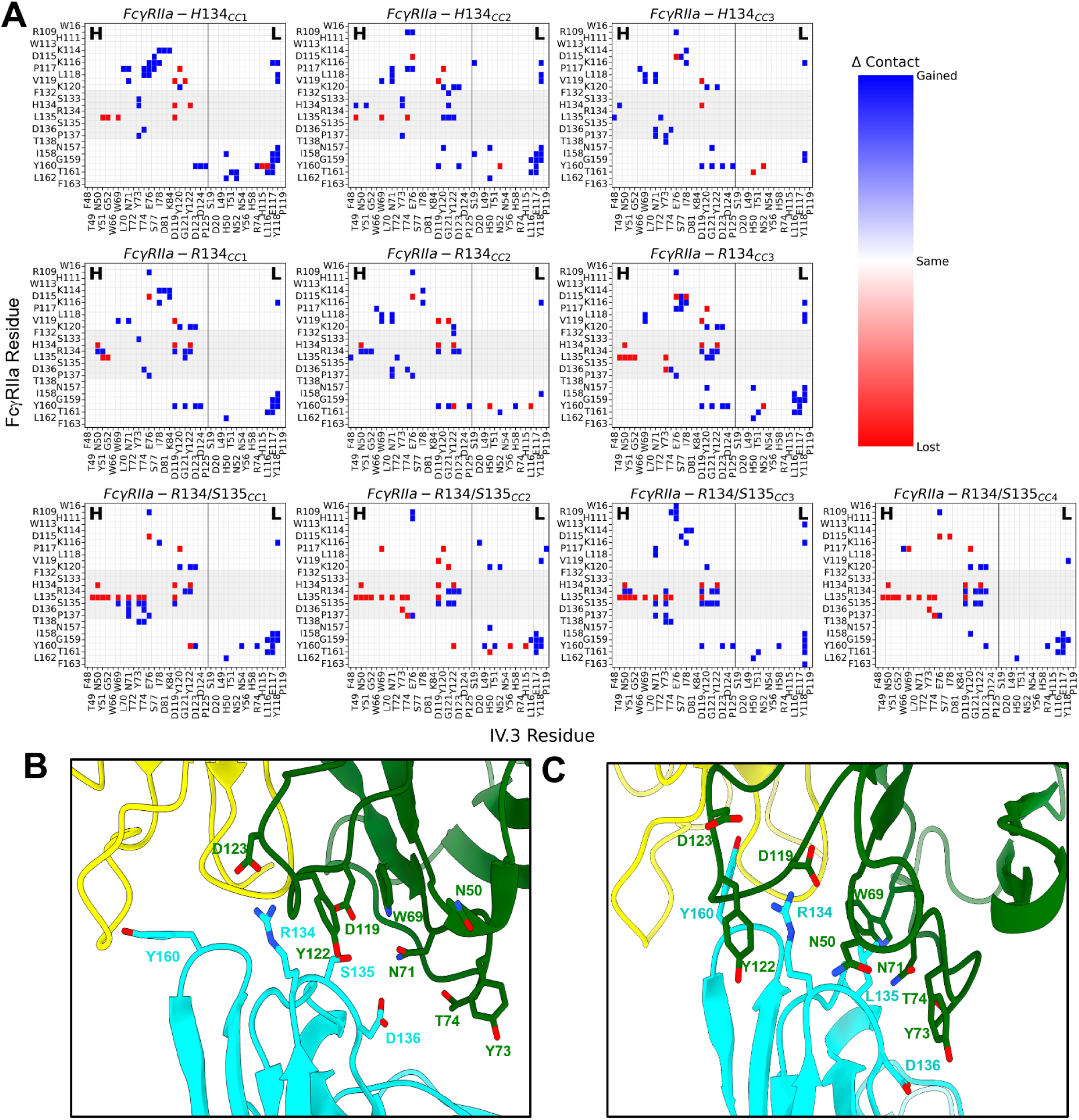
Clustering analysis of FcγRIIa dynamics. **(A)** Residue-wise contact–map divergence of cluster–center conformations for each simulated FcγRIIa system relative to the cryo–EM structure. Red indicates loss of contacts, white indicates no change, and blue denotes gain of a contact compared with the cryo-EM reference. The vertical black line separates the IV.3 heavy and light chains. The gray–shaded region highlights the FcγRIIa 132–137 epitope. The FcγRIIa R134/S135 construct exhibits a significant net loss of contacts across all cluster centers, particularly near residues W69 and Y73. Net contact changes per epitope residue are summarized in Supplementary Table 4; **(B)** Cluster–center #2 representative conformation of the FcγRIIa R134/S135 system showing that IV.3 heavy chain Y122 occludes R134, thereby preventing salt–bridge formation with IV.3 heavy chain D119 and instead promoting a transient interaction with IV.3 heavy chain D123, while FcγRIIa S135 forms a polar interaction with IV.3 heavy chain D119. Divergent conformations of FcγRIIa Y160 and IV.3 heavy chain Y73 further indicate that several key specificity–determining interactions are not formed, including those involving IV.3 heavy–chain residues N71 and T74; **(C)** Cluster–center #2 representative conformation of the FcγRIIa R134 system revealing a flipped IV.3 heavy chain Y122 loop that encloses FcγRIIa R134. This conformation stabilizes a backbone arrangement consistent with the experimental cryo–EM structure of the FcγRIIa–H134 variant, properly positioning FcγRIIa Y160 and IV.3 heavy chain Y73, and allowing FcγRIIa R134 to form a stable salt bridge with IV.3 heavy chain D119.

To further dissect the identified covarying interaction networks at the FcγRIIa–IV.3 interface, we compared feature distributions across simulations and quantified their difference using the Jensen–Shannon divergence^19^ with a significance threshold of 0.07 nats (Fig. 2D). Intra–chain distances involving FcγRIIa residues 135 and V119 differed significantly across all three simulated systems, with S135 displaying increased density at shorter distances. Among the most pronounced inter–chain dynamic variations were those observed in the IV.3–bound FcγRIIa R134/S135 system, where S135 approached IV.3 D119 more closely, forming hydrogen bonds and van der Waals contacts (Fig. 2C; left panel), thereby disrupting characteristic H/R134–mediated interactions (Fig. 2C; right panel).

Contact maps of the simulated FcγRIIa–R134/S135 complex (Fig. 3A; Supplementary Table 4) reveal a marked loss of contacts involving S135 relative to the cryo–EM FcγRIIa–H134 reference, particularly with residues W69 and Y73, consistent with the interaction differences highlighted in Figs. 2C–2D. Representative cluster–center structures from the FcγRIIa R134/S135 simulations (e.g., see cluster 2 representative in Fig. 3B) show that the IV.3 heavy–chain residue Y122 positions itself between FcγRIIa R134 and IV.3 heavy–chain residue D119, preventing the possible formation of a salt bridge between R134 and D119. Instead, R134 transiently engages D123 in interactions, while S135 forms polar and van der Waals interactions with D119.

In the R134/S135 system, divergent conformations of FcγRIIa Y160 and IV.3 heavy–chain residue Y73 further indicate the loss of several key specificity–determining contacts involving these residues, as well as IV.3 heavy–chain residues N71 and T74 (Fig. 3B). By contrast, simulations of the R134 variant reveal interactions with IV.3 heavy–chain residues D119, G121, and Y122, consistent with R134 inserting into this subpocket. Cluster–center analysis of the FcγRIIa R134 system further identifies a flipped conformation of IV.3 heavy–chain Y122 that encloses FcγRIIa R134 (e.g., see cluster 2 representative in Fig. 3C). This conformation shows the lowest RMSD relative to the cryo–EM structure of the FcγRIIa–H134 system (1.65 Å), faithfully recapitulates the orientations of FcγRIIa Y160 and IV.3 heavy–chain Y73, and enables R134 to form a stable salt bridge with IV.3 heavy–chain D119.

Together, these results support a mechanism in which L135–mediated hydrophobic stabilization primes a loop rearrangement in IV.3 that promotes formation of R134–dependent electrostatic, polar, cation–π and van der Waals interactions essential for maintaining FcγRIIa specificity. Conversely, substitution of L135 with serine disrupts these key hydrophobic contacts and introduces alternative interactions that interfere with the H/R134 interaction network. Additionally, Y160 acts as an interaction hub in both FcγRIIa polymorphs, forming shared contacts with the IV.3 heavy-chain residue Y120 (Figs. 1 and 3C), as well as polymorph-specific interactions. Notably, Y160 engages Y122 during simulations of the FcγRIIa-H134 variant but not of the FcγRIIa-R134 variant (Fig. 2C), underscoring subtle yet functionally meaningful differences in their protein–protein interaction fingerprints. These dynamics–driven structural changes provide a molecular basis for the selective recognition of FcγRIIa over FcγRIIb by mAb IV.3.

## Discussion

The 3.5 Å cryo–EM structure of the FcγRIIa–IV.3 Fab complex presented here provides a high–resolution view of how mAb IV.3 engages FcγRIIa and reveals a plausible mechanism of IgG Fc binding inhibition through steric hindrance. Consistent with earlier mutagenesis and chimeric receptor studies by Ramsland and colleagues,^9^ residues within the 132–137 region of FcγRIIa are confirmed as key determinants of mAb IV.3 recognition. Within this region, H134, L135, and D136 form a dense interaction network with multiple residues in the IV.3 heavy chain, including D119, Y122, W69, N71, and Y73.

Importantly, our cryo–EM structure extends these earlier observations by revealing that two additional FcγRIIa loop regions, specifically residues 113–120 and 156–163, also contribute directly to the IV.3 binding interface. Notably, P117, V119, and Y160 make extensive contacts across the interface and act as key interaction hubs. P117 forms hydrophobic interactions with IV.3 heavy–chain residues W69, I78, and Y120, as well as IV.3 light–chain residue Y118, and additionally engages in VdW interactions with IV.3 heavy–chain residue I78 and light-chain residue Y118. V119 forms a hydrophobic interaction with IV.3 heavy–chain residue Y120 and a VdW interaction with IV.3 heavy–chain residue D119. Y160 forms: (i) hydrophobic interactions with IV.3 heavy–chain residue Y120 and light–chain residues H50, Y56, and H115, (ii) a hydrogen bond through its backbone carbonyl oxygen with the Nε2 atom of the IV.3 light chain H50 imidazole ring, (iii) VdW interactions with IV.3 light–chain residues H50, Y56, and H115, and (iv) face–to–face and π–π stacking interactions with Y56. These observations explain why earlier domain–swap experiments^9^ failed to fully map the IV.3 epitope. Although the corresponding amino acid to Y160 in the FcγεRI construct used for swapping is a glutamine, the engineered chimera substituted only residues 150–172, excluding the additional contributing regions, residues 113–120 and 133–137, indicating that Y160 alone is insufficient to support IV.3 binding. Likewise, a second chimera containing FcγRIIa Y160, but lacking H134 and L135 did not bind IV.3,^9^ further demonstrating that, despite its prominent role, Y160 requires cooperative interactions with neighboring residues to enable high–affinity recognition.

Our SPR measurements confirm previous reports that mAb IV.3 binds both FcγRIIa polymorphic variants, FcγRIIa–H134 and FcγRIIa–R134, with subnanomolar affinities,^7^ while exhibiting negligible affinity for FcγRIIb.^2^ Alchemical free energy calculations further support this specificity: the double H134R/L135S transformation in FcγRIIa, which mimics the FcγRIIb epitope, results in a pronounced loss of binding free energy, consistent with the experimental specificity profile. In contrast, simulations of the single H134R and L135S perturbations indicated only modest changes in FcγRIIa–IV.3 Fab binding free energy, with values falling within the typical uncertainty range of free energy estimation methods (1.2–2.0 kcal·mol⁻¹).^15, 20, 21^

A broader limitation of alchemical free energy methods is their difficulty in capturing slow conformational transitions that can substantially influence the endpoint states, such as, for instance, loop rearrangements that typically occur on microsecond or longer timescales.^22^ Although enhanced sampling approaches can accelerate conformational sampling, sampling convergence across coupled alchemical and conformational degrees of freedom remains computationally demanding. To evaluate whether initiating our free energy calculations from the cryo–EM structure of the FcγRIIa–H134 variant bound to IV.3 Fab, where H134 forms native contacts with IV.3 heavy–chain residue D119, might bias the results due to limited conformational sampling, we performed MD simulations of the H134, R134 and R134/S135 variants of the FcγRIIa–IV.3 Fab complex. TICA projections of the resulting trajectories revealed distinct conformational dynamics among the three simulated IV.3–bound FcγRIIa complexes, motivating a closer examination of these dynamics to gain deeper insight into the determinants of IV.3 binding specificity.

Time–lagged covariance network analysis of dynamic features identified FcγRIIa residues 134 and 135 as key dynamic hubs that coordinate correlated motions of adjacent loops through networks involving IV.3 heavy–chain residues N71, T72, Y73, T74, D119, G121, and Y122, as well as FcγRIIa residues K120 and V119. Y160 was also implicated in this network through covariation with IV.3 heavy–chain residues G121 and Y120, as well as IV.3 light–chain residue H50. The R134 variant stabilizes this network through cation–π, salt–bridge, polar, and van der Waals interactions, whereas L135 contributes hydrophobic packing that positions R134 and Y160 optimally for Fab engagement. In the FcγRIIb–like FcγRIIa–R134/S135 complex, disruption of these interactions reconfigures the conformational ensemble, weakening key hydrophobic and aromatic contacts and promoting occlusion of R134 from interaction with D119 through Y122 repositioning, thereby reducing binding affinity. Collectively, these results underscore the interplay between local side–chain chemistry and collective loop dynamics in defining FcγRIIa–antibody specificity.

Beyond elucidating the structural basis of mAb IV.3 selective recognition of FcγRIIa over FcγRIIb, these results have direct translational implications. FcγRIIa–mediated immune complex binding contributes to a range of inflammatory and autoimmune pathologies (reviewed in references^23, 24^). The atomic–level description of the IV.3 epitope and mechanistic basis for FcγRIIa specificity provide a valuable template for designing next–generation inhibitors that mimic critical receptor–antibody interactions while minimizing Fc–mediated effector functions. Indeed, the previously cited humanized version of mAb IV.3 in which mutations were introduced to reduce Fc–mediated binding to Fc receptors^7^ was reported to: (i) protect against immune thrombocytopenia, neutrophil infiltration in response to anti–glomerular basement membrane–induced nephritis, and anti–collagen antibody–induced arthritis in murine models; (ii) have a favorable safety profile in non–human primates after three months of therapy; and (iii) have favorable pharmacokinetics and pharmacodynamics. A Phase 1 study with this antibody was started in 2018 but is currently listed as terminated on ClinicalTrials.gov.^25^ The present structure–function framework offers new opportunities for rational engineering of FcγRIIa antagonists, including small molecules or engineered biologics that selectively block pathogenic FcγRIIa signaling without interfering with FcγRIIb’s regulatory functions. Such therapeutics hold the promise of being antithrombotic in disorders characterized by thromboinflammation induced by immune complexes, without increasing the risk of bleeding, a goal that has remained elusive to date.

## Methods

### Binding affinity and kinetics by surface plasmon resonance

Binding kinetics of the murine mAb IV.3 Fab to human FcγRIIb, FcγRIIa–H134, and FcγIIa–R134 were determined using a Biacore T200 SPR system (Cytiva Life Sciences) at 25°C in HBS–EP+ buffer (pH 7.4), as previously described.^11^ To prevent Fc–FcγR interactions, the IV.3 mAb was recombinantly expressed in the human IgG1 backbone (as previously described^11^), encompassing mutations at the Fc domain (G236R/L328R) to abrogate FcγR binding. mAb IV.3 at a concentration of 10 µg/ml was immobilized for 60 s at 20 µl/min on a Series S protein G sensor chip (Cytiva) to achieve a target density of 2,000 response units (RU). Soluble human Fcγ receptor ectodomains (FcγRIIb, FcγRIIa–H134, and FcγRIIa–R134) were each flowed at 20 µl/min for 60 s at the indicated concentrations, followed by a 600 s dissociation phase. After each binding cycle, the sensor chip was regenerated with glycine–HCl (pH 1.5) for 30 s at 50 µl/min. Biacore evaluation software 3.2.1 was used to calculate kinetic constants. Sensorgrams were fit using the 1:1 Langmuir binding model (Supplementary Fig. 7) and affinity constants (K_D_) were calculated.

### Complex formation and negative–stain electron microscopy

Lyophilized human FcγRIIa extracellular domain (Met1–Ile218) with a C–terminal polyhistidine tag, purchased from Sino Biological Inc., was dissolved in sterile PBS (pH 7.4) to 0.2 mg/ml and mixed with IV.3 Fab in PBS buffer, pH 7.4 at a 1:2 molar ratio. After 1 h at 4°C, a 3.5 μL aliquot of the solution (∼14 µg) was applied to a glow–discharged thin carbon film that had been evaporated onto a plastic–covered copper grid. After 30s, the grid was blotted, washed twice with deionized water, and stained with 0.7% (w/v) uranyl formate. Grids were imaged with a Philips CM10 electron microscope operated at an acceleration voltage of 80 kV using a defocus of about −1.5 μm and a nominal magnification of 52,000×, yielding a calibrated pixel size on the specimen level of 1.97 Å. A total of 47 images were collected with an AMT 3k × 5k ActiveVu CCD camera.

Particles were automatically picked with the Blob Picker tool in CryoSPARC,^26^ which was used for all further image processing. An initial set of 28,424 particles was subjected to 2D classification. Class averages displaying clear structural features of the FcγRIIa–IV.3 Fab complex were then used as a template for a second round of particle picking, yielding 56,510 particles. Classes with uninformative or poorly defined features were discarded, and the remaining particles underwent four additional iterative rounds of 2D classification. The final dataset contained 16,216 particles, which were extracted into 140×140–pixel images, centered, normalized, and subjected to 2D classification into 100 groups, yielding the 2D class averages shown in Supplementary Fig. 1.

### Cryo–EM sample preparation and data collection

A 3.5–µL aliquot of the FcγRIIa–IV.3 Fab mixture used for negative–stain EM analysis was applied to a freshly glow–discharged 400–mesh graphene oxide copper grid (GO, R1.2/1.3). Grids were blotted with filter paper for 1.5 s under 100% relative humidity and plunge–frozen in liquid ethane (Vitrobot; FEI). Cryo–EM data sets were collected on a Titan Krios microscope (FEI) at 300 kV. Movies comprising 40 frames (0.05 s per frame; total electron dose of 54 e^−^/Å2) were recorded in super–resolution mode on a K3 direct detector with a defocus range of –1.2 to –2.5 μm. Movies were acquired with SerialEM^27^ at a nominal magnification of 81,000×, corresponding to a calibrated pixel size of 0.86 Å. Cryo–EM data collection statistics are provided in Table S1.

### Cryo–EM image processing

A total of 8,898 image stacks were motion–corrected, dose–weighted, summed, and binned over 2×2 pixels using MotionCor2.^28^ The contrast transfer function (CTF) parameters were determined with cryoSPARC,^26^ which was also used for all subsequent image processing steps. During exposure curation, 1,267 of the 8,898 micrographs were removed, primarily based on a CTF fit resolution threshold below 5 Å. Blob picking was used to auto–pick particles from 2,000 out of the remaining 7,631 micrographs without templates, yielding 42,902 high–quality particles that were used to train a Topaz picking model.^29^ Using this model, 2,705,081 particles were automatically picked and extracted into 400×400–pixel images, which were binned over 4×4 pixels, and subjected to 2D classification into 100 classes. Particles from the 14 classes showing the clearest features of the FcγRIIa–IV.3 Fab complex (450,922 particles) were combined and used to generate four *ab–initio* models. Two of these models displayed characteristic features of the FcγRIIa–IV.3 Fab complex, and one of them (*ab–initio* model 1) was selected as a reference for subsequent 3D classification steps.

In parallel, classes from the initial 2D classification that produced uninformative averages (representing ice contaminations or edge artifacts) were removed from the dataset. The remaining particles were subjected to another round of 2D classification, yielding 93 class averages that showed well–defined views of the FcγRIIa–IV.3 Fab complex. The 2,102,469 particles contributing to these classes were combined and used for 3D classification into 8 classes, using as references four copies of *ab–initio* model 1 and four noise maps. Particles belonging to the four classes that had *ab–initio* model 1 as the reference (1,148,107 particles) were combined and subjected to three additional rounds of 3D classification using as references four copies of *ab–initio* model 1 and four noise maps. After removing particles assigned to the four noise maps, the remaining 392,060 particles were combined, re–centered, re–extracted into 400×400–pixel images, and refined using non–uniform refinement, resulting in a final reconstruction at 3.5–Å resolution. Local resolution estimates of the final map was calculated in cryoSPARC.

### Model building and refinement

To build the model of the FcγRIIa–IV.3 Fab complex, the crystal structure of the FcγRIIa ectodomain (Protein Data Bank [PDB] ID: 1H9V) and the IV.3 Fab model predicted by AlphaFold2 were rigid–body fitted into the 3.5 Å–resolution cryo–EM density map using Chimera.^30^ After residues were fitted into the map, some were manually rebuilt and adjusted in Coot.^31^ The resulting model, comprising the FcγRIIa ectodomain (residues 4 to 175) and the IV.3 Fab heavy–(residues 19–238) and light– (residues 19–238) chains, was refined through iterative cycles of real–space refinement in Phenix,^32^ followed by minor manual adjustments guided by Ramachandran, rotamer, and clashscore restraints. Final model geometry was assessed in the context of the available map resolution using MolProbity on the official PDB validation server. Refinement statistics for the final FcγRIIa–IV.3 Fab model are summarized in Supplementary Table 1.

### Sequence alignment

Sequence alignment of FcγRIIa and FcγRIIb was performed using the EMBL–EBL Dispatcher sequence analysis tool.^33^

### Alchemical free energy calculations of complex and variants

The expanded ensemble (EE) method^15^ was applied to compute alchemical free energies for FcγRIIa both in solvent and in complex with the mAb IV.3 Fab, thereby determining sequence–dependent free energy changes in thermostability and binding affinity. The EE approach samples all thermodynamic intermediates within a single simulation, accelerating alchemical ladder sampling by iteratively biasing under–sampled λ–states using the Wang–Landau flat–histogram algorithm.^34, 35^ Once histogram flatness is reached, the Wang–Landau bias increment is reduced, and sampling continues with this smaller bias increment. When the bias increment becomes sufficiently small, the free–energy difference between end states can be directly calculated as a time–dependent mean of the accumulated bias. As with other alchemical approaches, however, the EE method faces challenges when transformations involve charge changes, slow conformational dynamics, and large alchemical perturbations (≳25 alchemical atoms).^15^ Additionally, the GROMACS implementation of EE requires use of the velocity–Verlet integrator,^36^ restricting force evaluations to CPUs rather than GPUs, reducing computational efficiency.

Starting from an earlier refinement of the deposited cryo–EM structure of the FcγRIIa–IV.3 Fab complex, alchemical topologies and structure files were generated using PMX^37^ and GROMACS tools^38^ using the amber14sbmut forcefield.^37, 39^ Specifically, the following transformations were performed: the single L135S and H134R perturbations, and the combined H134R/L135S transformation. GROMACS version 2025.1^38^ compiled with CUDA 12.4^40^ was used for system preparation and simulation. For the L135S and H134R single perturbations, the L2S and X2R (HIE alchemical His) residues were used, respectively. For the H134R/L135S double transformation, the X2R alchemical residue was used, and L135 was mutated to serine in ChimeraX^41^ using the Dunbrack rotamer library (which is also used in PMX to assign alchemical sidechain rotamers)^42^

Each system was placed in a cubic box with a 1 nm box edge buffer (complex: 13.3 nm^3^ and non–complex: 8.3 nm^3^) and solvated using TIP3P^43^ water and Joung–Cheatham ion parameters.^44, 45^ In total, the simulated FcγRIIa–IV.3 Fab complex systems had 235,026 (H134R), 235,003 (L135S), and 235,033 (H134R/L135S) atoms respectively, and the FcγRIIa–only systems had 56,835, 56,848, and 56,830 atoms respectively (Supplementary Table 6). Sodium and chloride ions were added, replacing randomly selected water molecules to neutralize the protein charge and achieve a 0.15 M salt concentration. To conserve charge during charge changing (X2R) alchemical transformations, a single sodium ion was replaced with an alchemical NAZ atom, restrained to a corner of the box with a force–constant of 1000 kj mol^-1^ nm^-1^. This atom was gradually transformed from interacting to non–interacting to preserve overall charge neutrality.^46^

Each system was energy minimized using a steepest–descent algorithm for 10,000 steps and equilibrated to 300 K for 100 ps each under NPT and NVT ensembles. For production EE simulations, hydrogens were constrained using LInear Constraint Solver (LINCS),^47^ Particle Mesh Ewald (PME) electrostatics^48^, 0.9 nm van der Waals (vdW) cutoffs, and long–range dispersion corrections. For the X2R mutations, hydrogen mass repartitioning (HMR) was applied by setting the minimum hydrogen mass to 3 amu, enabling a 4 fs integration timestep. For the L2S system, a 2 fs timestep was used since HMR cannot be applied to alchemically transformed hydrogens.

Simulations used an H100 GPU for bonded, non–bonded, and PME calculations, while force updates were done using 24 Intel Xeon Platinum 8568Y+ processor cores. The GROMACS implementation of EE^49^ was used with 11 (L2R) or 109 (X2R) intermediates along the alchemical transformation. Soft–core potentials were used with sc–α = 0.5, sc–power = 1, and sc–σ = 0.3. Exchanges were proposed/accepted every 1 ps according to a Metropolis criterion.^50^ The initial Wang−Landau increment^34, 35^ was set to δ = 10 kBT. The sampling histogram was deemed sufficiently flat when all histograms counts h_i_ came within some threshold of the mean,

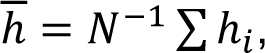

such that *min(h_i_/<u>h</u>,<u>h</u>/h_i_) > η* for all i, where η is set to 0.8. Upon reaching this criterion, the histogram counts were reset to zero and the Wang−Landau increment was scaled by α = 0.5. An initial 20 ns EE^45^ test simulation was carried out using the velocity–Verlet integrator^36^ in the NVT ensemble to evaluate sampling efficiency. After this test simulation, the *pylambdaopt* algorithm^46^ was used to optimize the number and spacing of λ intermediates for the X2R residues, yielding an optimal λ schedule of 109 alchemical intermediates (versus the initial 11; see initial and final lambda values in Supplementary Table 4), thereby addressing sampling inefficiencies associated with the arginine alchemical transformation.^15^ We report the final free energies as time–dependent means obtained after the bias increment has reached a sufficiently small log_10_ value [L2S: (complex and non–complex: –4); X2R: (complex:–1.5, non–complex:–2]), and we estimate the uncertainties by propagating the standard error of the mean (SEM) from both legs of the alchemical transformation.

### Interfacial dynamics in the complex and selected mutants

Standard molecular dynamics simulations were performed on an earlier refinement of the deposited cryo–EM structure of the FcγRIIa–IV.3 Fab complex, as well as on variant models in which H134 was substituted with arginine (R134) or both H134 and L135 were substituted with arginine and serine (R134/S135). System setup details are provided in Supplementary Table 7. The single (R134) and double (R134/S135) mutations were introduced into the cryo–EM FcγRIIa–IV.3 Fab structure in ChimeraX using the Dunbrack rotomer library.^41, 42^ System topologies were generated with the amber14sb forcefield^39^ and simulations were performed using GROMACS 2025.1 on an H100 GPU. Each system was energy minimized using the steepest–descent algorithm for 10,000 steps and equilibrated at 300 K for 100 ps in the NVT and NPT ensembles. For production simulations, hydrogens were constrained using LINCS, PME electrostatics, 0.9 nm vdW cutoffs, and long–range dispersion corrections. HMR was used as described above. Production NPT simulations were run in triplicate for a total simulation time of ∼1.9 μs for each system using a leap–frog integrator.

Trajectory analyses were performed using MDTraj,^51^ Deeptime,^52^ MDAnalysis,^53^ and ProLIF.^14^ Trajectories were featurized using inter-residue Cα_i_–Cα_j_ and Cβ_i_–Cβ_j_ distances, as well as Cα_i_–Cβ_i_–Cβ_j_–Cα_j_ torsion angles computed between residues forming heavy atom contacts (≤4.5 Å) with Y160, K120, and residues at positions 134 or 135. These features were analyzed via tICA,^16, 17^ using commute–distance mapping and a 20 ns lag time. The resulting time–lagged covariance matrix yielded a weighted feature covariance network, from which a maximal-weight connected feature path was identified using a greedy heuristic algorithm^18^ (Eq. 1). Specifically, let *G* = (*V*, *E*) be an undirected graph in which nodes (*V*) represent FcγRIIa-IV.3 interfacial residues and edges (*E*) are weighted by the corresponding elements of the time-lagged covariance matrix, *w*: *E* → ℝ, where *w*(*u*, *v*). A path *P* = (*v*_0_, …, *v*_*k*_) is defined as simple if all nodes are distinct and consecutive nodes are adjacent.

The path construction is initialized from the edge with maximum weight,

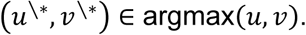

At each iteration, the current path endpoints *v*_0_ and *v*_*k*_ are extended by considering all unused neighboring nodes,

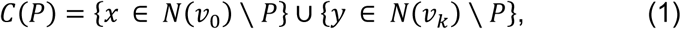

with each candidate scored by the weight of its incident edge (i.e., the residue-residue covariance),

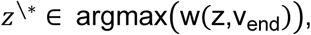

The candidate associated with the largest edge weight is appended to the corresponding end of the path, and nodes already present in *P* are excluded to preserve path simplicity. This procedure is repeated until no further extensions are possible.

Finally, differences in feature distributions between systems were quantified using the Jensen–Shannon divergence,^19^ with a threshold of 0.07 nats indicating significant divergence:

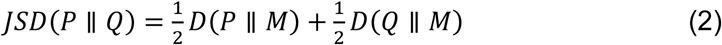

where *M(P ∥ Q) = ½(P + Q)*

## Acknowledgements

This work was supported by NIH grant HL019278 (B.C. and M.F.) Computational work was supported in part through the computational resources and staff expertise provided by Scientific Computing at the Icahn School of Medicine at Mount Sinai and supported by the Clinical and Translational Science Awards (CTSA) grant UL1TR004419 from the National Center for Advancing Translational Sciences. Research reported in this paper was also supported by the Office of Research Infrastructure of the National Institutes of Health under award number S10OD026880 and S10OD030463. The content is solely the responsibility of the authors and does not necessarily represent the official views of the National Institutes of Health.

## Author Contributions

J.W. designed cryo-EM experiments, collected data, refined structures, and contributed to writing the manuscript. D.N. designed, performed, and analyzed computational simulations and contributed to writing the manuscript. J.L. assisted in the conduct of the cryo-EM studies. E.G.N. performed the SPR experiments under the supervision of S.B., who also contributed to writing the manuscript. M.F. supervised the computational work, analyzed related data, and wrote the manuscript. B.S.C. supervised all experimental work, analyzed related data, and wrote the manuscript.

## Data Availability

The cryo–EM density map and atomic coordinates for the FcγRIIa–IV.3 Fab complex have been deposited in the Electron Microscopy Data Bank (accession codes EMD–75133) and the Protein Data Bank (accession code 10FJ), respectively.

## References

1. McKenzie SE, Goldfinger LE, Michael JV. Platelet FcγRIIA Receptor. In: Platelet Physiology I. Platelets in Thrombotic and Non-Thrombotic Disorders. (ed Gresele P, López, J.A., Angiolillo, D.J., Page, C.P.). Springer, Cham. (2025).

2. Rosenfeld SI, Looney RJ, Leddy JP, Phipps DC, Abraham GN, Anderson CL. Human platelet Fc receptor for immunoglobulin G. Identification as a 40,000-molecular-weight membrane protein shared by monocytes. J Clin Invest 76, 2317–2322 (1985).

3. Gao C, et al. Eptifibatide-induced thrombocytopenia and thrombosis in humans require FcgammaRIIa and the integrin beta3 cytoplasmic domain. J Clin Invest 119, 504–511 (2009).

4. Leung HHL, et al. Antithrombotic efficacy and bleeding risks of vaccine-induced immune thrombotic thrombocytopenia treatments. Blood Adv 8, 5744–5752 (2024).

5. Martins-Goncalves R, et al. Platelet-neutrophil aggregate formation induces NLRP3 inflammasome activation in vaccine-induced thrombotic thrombocytopenia. J Thromb Haemost 23, 1034–1042 (2025).

6. Palankar R, Kohler TP, Krauel K, Wesche J, Hammerschmidt S, Greinacher A. Platelets kill bacteria by bridging innate and adaptive immunity via platelet factor 4 and FcgammaRIIA. J Thromb Haemost 16, 1187–1197 (2018).

7. Chen B, et al. Humanised effector-null FcgammaRIIA antibody inhibits immune complex-mediated proinflammatory responses. Ann Rheum Dis 78, 228–237 (2019).

8. Powell MS, et al. Biochemical analysis and crystallisation of Fc gamma RIIa, the low affinity receptor for IgG. Immunol Lett 68, 17–23 (1999).

9. Ramsland PA, et al. Structural basis for Fc gammaRIIa recognition of human IgG and formation of inflammatory signaling complexes. J Immunol 187, 3208–3217 (2011).

10. Morganelli PM, Groveman DS, Pfeiffer JR. Evidence that human Fc gamma receptor IIA (CD32) subtypes are not receptors for oxidized LDL. Arterioscler Thromb Vasc Biol 17, 3248–3254 (1997).

11. Bournazos S, Corti D, Virgin HW, Ravetch JV. Fc-optimized antibodies elicit CD8 immunity to viral respiratory infection. Nature 588, 485–490 (2020).

12. Veri MC, et al. Monoclonal antibodies capable of discriminating the human inhibitory Fcgamma-receptor IIB (CD32B) from the activating Fcgamma-receptor IIA (CD32A): biochemical, biological and functional characterization. Immunology 121, 392–404 (2007).

13. Jones AT, Marino AE, Martynyuk T, Bournazos S, Ravetch JV. The anti-inflammatory activity of IgG is enhanced by co-engagement of type I and II Fc receptors. Science 390, eadv2927 (2025).

14. Bouysset C, Fiorucci S. ProLIF: a library to encode molecular interactions as fingerprints. J Cheminform 13, 72 (2021).

15. Novack D, Zhang S, Voelz VA. Massively Parallel Free Energy Calculations for In Silico Affinity Maturation of Designed Miniproteins. J Chem Theory Comput 21, 8034–8050 (2025).

16. Molgedey L, Schuster HG. Separation of a mixture of independent signals using time delayed correlations. PhysRev Lett 72, 3634 (1994).

17. Naritomi Y, Fuchigami S. Slow dynamics of a protein backbone in molecular dynamics simulation revealed by time-structure based independent component analysis. J Chem Phys 139, 215102 (2013).

18. Cormen TH, Leiserson CE, Rivest RL, Stein C. Introduction to Algorithms, 4th ed.). MIT Press (2022).

19. Lin J. Divergence measures based on the Shannon entropy. IEEE Transactions on Information Theory 37, 145–151 (1991).

20. Aldeghi M, Gapsys V, de Groot BL. Accurate Estimation of Ligand Binding Affinity Changes upon Protein Mutation. ACS Central Science 4, 1708–1718 (2018).

21. Singh S, et al. Prospective Evaluation of Structure-Based Simulations Reveal Their Ability to Predict the Impact of Kinase Mutations on Inhibitor Binding. J Phys Chem B 129, 2882–2902 (2025).

22. Shaw DE, et al. Atomic-level characterization of the structural dynamics of proteins. Science 330, 341–346 (2010).

23. Muller L, Dabbiru VAS, Schonborn L, Greinacher A. Therapeutic strategies in FcgammaIIA receptor-dependent thrombosis and thromboinflammation as seen in heparin-induced thrombocytopenia (HIT) and vaccine-induced immune thrombocytopenia and thrombosis (VITT). Expert Opin Pharmacother 25, 281–294 (2024).

24. Patel P, Michael JV, Naik UP, McKenzie SE. Platelet FcgammaRIIA in immunity and thrombosis: Adaptive immunothrombosis. J Thromb Haemost 19, 1149–1160 (2021).

25. https://clinicaltrials.gov/search?intr=vib9600.).

26. Punjani A, Rubinstein JL, Fleet DJ, Brubaker MA. cryoSPARC: algorithms for rapid unsupervised cryo-EM structure determination. Nat Methods 14, 290–296 (2017).

27. Mastronarde DN. Automated electron microscope tomography using robust prediction of specimen movements. J Struct Biol 152, 36–51 (2005).

28. Zheng SQ, Palovcak E, Armache JP, Verba KA, Cheng Y, Agard DA. MotionCor2: anisotropic correction of beam-induced motion for improved cryo-electron microscopy. Nat Methods 14, 331–332 (2017).

29. Bepler T, et al. Positive-unlabeled convolutional neural networks for particle picking in cryo-electron micrographs. Nat Methods 16, 1153–1160 (2019).

30. Pettersen EF, et al. UCSF ChimeraX: Structure visualization for researchers, educators, and developers. Protein Sci 30, 70–82 (2021).

31. Emsley P, Lohkamp B, Scott WG, Cowtan K. Features and development of Coot. Acta Crystallogr D Biol Crystallogr 66, 486–501 (2010).

32. Afonine PV, et al. Towards automated crystallographic structure refinement with phenix.refine. Acta Crystallogr D Biol Crystallogr 68, 352–367 (2012).

33. Madeira F, et al. The EMBL-EBI Job Dispatcher sequence analysis tools framework in 2024. Nucleic Acids Res 52, W521–W525 (2024).

34. Wang F, Landau DP. Efficient, multiple-range random walk algorithm to calculate the density of states. Phys Rev Lett 86, 2050–2053 (2001).

35. Wang F, Landau DP. Determining the density of states for classical statistical models: a random walk algorithm to produce a flat histogram. Phys Rev E Stat Nonlin Soft Matter Phys 64, 056101 (2001).

36. Swope WC, Andersen HC, Berens PH, Wilson KR. A computer simulation method for the calculation of equilibrium constants for the formation of physical clusters of molecules: Application to small water clusters. The Journal of Chemical Physics 76, 637–649 (1982).

37. Gapsys V, de Groot BL. pmx Webserver: A User Friendly Interface for Alchemistry. J Chem Inf Model 57, 109–114 (2017).

38. Abraham M, et al. GROMACS 2025.1 Manual.) (2025).

39. Maier JA, Martinez C, Kasavajhala K, Wickstrom L, Hauser KE, Simmerling C. ff14SB: Improving the Accuracy of Protein Side Chain and Backbone Parameters from ff99SB. J Chem Theory Comput 11, 3696–3713 (2015).

40. CUDA Toolkit Documentation 12.4..) (2025).

41. Meng EC, et al. UCSF ChimeraX: Tools for structure building and analysis. Protein Sci 32, e4792 (2023).

42. Shapovalov MV, Dunbrack RL, Jr. A smoothed backbone-dependent rotamer library for proteins derived from adaptive kernel density estimates and regressions. Structure 19, 844–858 (2011).

43. Jorgensen WL, Chandrasekhar J, Madura JD, Impey RW, Klein ML. Comparison of Simple Potential Functions for Simulating Liquid Water. J Chem Phys 79, 926–935 (1983).

44. Joung IS, Cheatham TE, 3rd. Determination of alkali and halide monovalent ion parameters for use in explicitly solvated biomolecular simulations. J Phys Chem B 112, 9020–9041 (2008).

45. Joung IS, Cheatham TE, 3rd. Molecular dynamics simulations of the dynamic and energetic properties of alkali and halide ions using water-model-specific ion parameters. J Phys Chem B 113, 13279–13290 (2009).

46. Novack D, Raddi RM, Zhang S, Hurley MFD, Voelz VA. Simple Method to Optimize the Spacing and Number of Alchemical Intermediates in Expanded Ensemble Free Energy Calculations. J Chem Inf Model 65, 6089–6101 (2025).

47. Hess B, Bekker H, Berendsen HJC, Fraaije JGEM. LINCS: A linear constraint solver for molecular simulations. Journal of Computational Chemistry 18, 1463–1472 (1998).

48. Darden T, York D, Pedersen L. Particle mesh Ewald: An N⋅log(N) method for Ewald sums in large systems. The Journal of Chemical Physics 98, 10089–10092 (1993).

49. Chodera JD, Shirts MR. Replica exchange and expanded ensemble simulations as Gibbs sampling: simple improvements for enhanced mixing. J Chem Phys 135, 194110 (2011).

50. Metropolis N, Risenbluth AW, Rosenbluth MN, Teller AH, Teller E. Equation of State Calculations by Fast Computing Machines. The Journal of Chemical Physics 21, 1087–1092 (1953).

51. McGibbon RT, et al. MDTraj: A Modern Open Library for the Analysis of Molecular Dynamics Trajectories. Biophys J 109, 1528–1532 (2015).

52. Hoffmann M, et al. Deeptime: A Python Library for Machine Learning Dynamical Models from Time Series Data. Mach Learn Sci Technol 3, 015009 (2021).

53. Naughton FB, et al. MDAnalysis 2.0 and beyond: Fast and Interoperable, Community Driven Simulation Analysis. Biophys J 121, 272a–273a (2022).

