## Supplementary Material for "Mechanistic Basis for the Selective Recognition of the Fcγ Receptor IIa by Monoclonal Antibody IV.3"

**List of Supplementary Materials:**

**Supplementary Fig. S1.** Negative–stain EM analysis of the FcγRIIa ectodomain in complex with mAb IV.3 Fab.

**Supplementary Fig. S2.** Cryo–EM data–processing workflow for the FcγRIIa–IV.3 Fab complex.

**Supplementary Fig. S3.** Cryo–EM reconstruction quality assessment for the FcγRIIa–IV.3 Fab complex.

**Supplementary Fig. S4.** Sequence comparison between FcγRIIa and FcγRIIb.

**Supplementary Fig. S5.** Comparison of the FcγRIIa–IV.3 Fab cryo–EM structure with the previously published crystal structure of the FcγRIIa–R134 variant bound to the Fc domain of human IgG1.

**Supplementary Fig. S6.** Theoretical framework for alchemical free energy calculations.

**Supplementary Fig. S7.** Surface plasmon resonance analysis of FcγRII binding to mAb IV.3.

**Supplementary Table S1.** Cryo–EM data collection, refinement and validation statistics.

**Supplementary Table S2.** FcgRIIa–IV.3 and FcgRIIa–IgG1 (3RY6) protein–protein interaction fingerprints.

**Supplementary Table S3.** Mutational free–energy estimates of the FcγRIIa–IV.3 Fab complex.

**Supplementary Table S4.** Net delta contacts per FcγRIIa epitope residue.

**Supplementary Table S5.** Initial and final lambdas.

**Supplementary Table S6**. MD simulation setup

**Supplementary Table S7**. Alchemical Expanded Ensemble simulation setup

**Supplementary Movie 1.** mAb IV.3 Fab Heavy Chain Loop 1 Rearrangement Around FcγRIIa–R134.

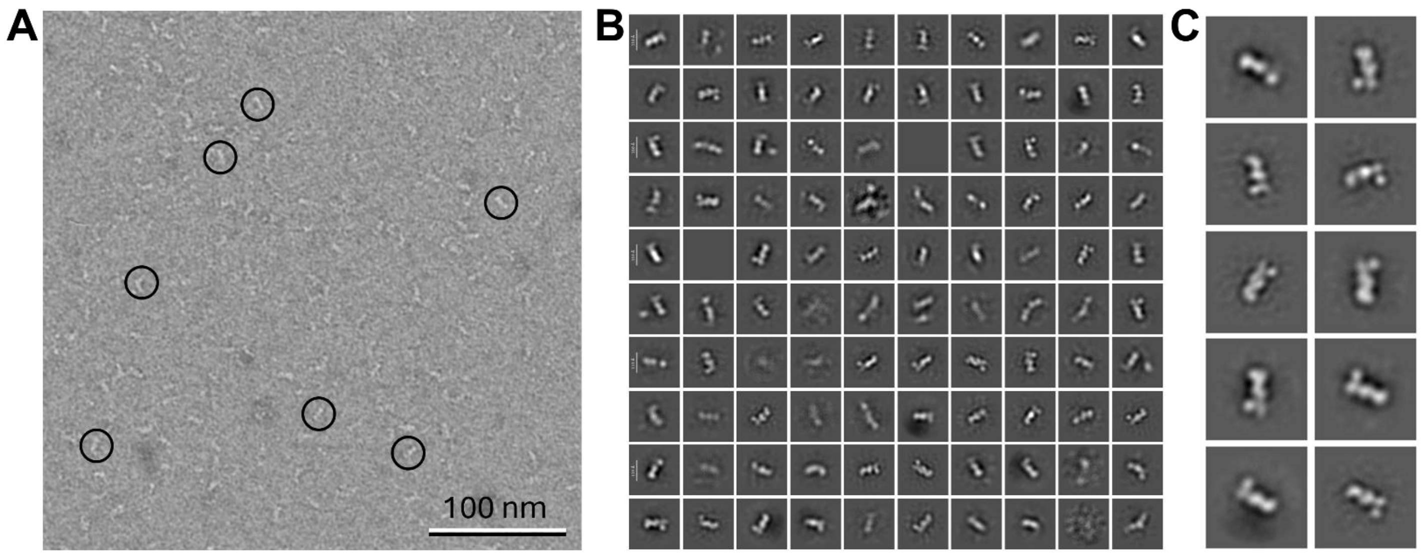


**Supplementary Fig. S1. Negative–stain EM analysis of the FcγRIIa ectodomain in complex with mAb IV.3 Fab. (A)** Representative raw micrograph of negatively stained FcγRIIa–IV.3 Fab complexes. Individual particles are indicated by circles. **(B)** A set of 100 two–dimensional (2D) class averages of the FcγRIIa–IV.3 Fab complex. **(C)** Representative class averages illustrating the complex in different orientations. The scale bar corresponds to 100 nm; the box size for the averages is 27.6 nm.

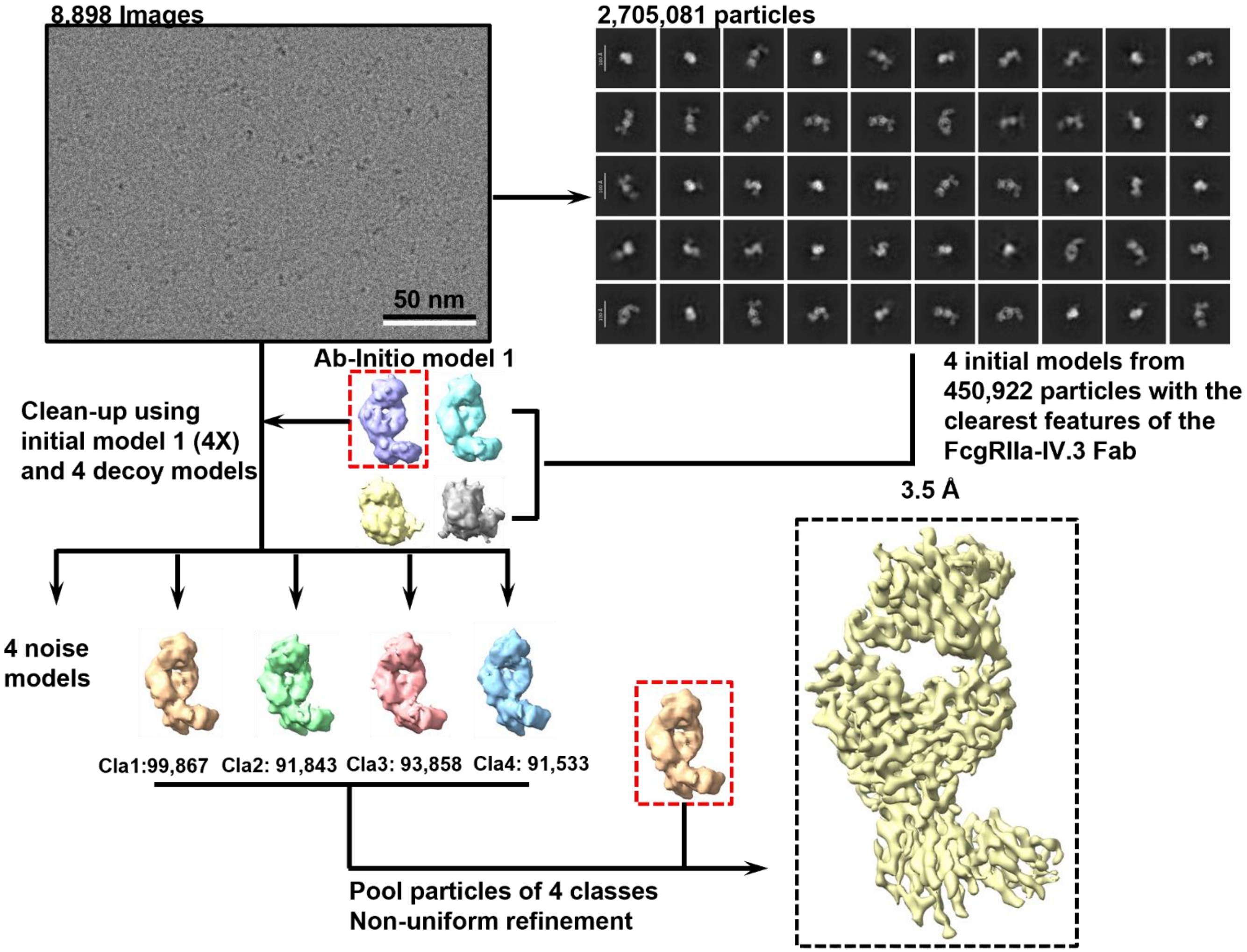


**Supplementary Fig. S2. Cryo–EM data–processing workflow for the FcγRIIa–IV.3 Fab complex.** Flowchart summarizing the cryo–EM processing pipeline for the FcγRIIa–IV.3 Fab complex in CryoSPARC. Additional methodological details are provided in the *Methods* section. The final reconstruction yielded density maps of the FcγRIIa–IV.3 Fab complex at 3.5–Å resolution. Scale bar, 50 nm.

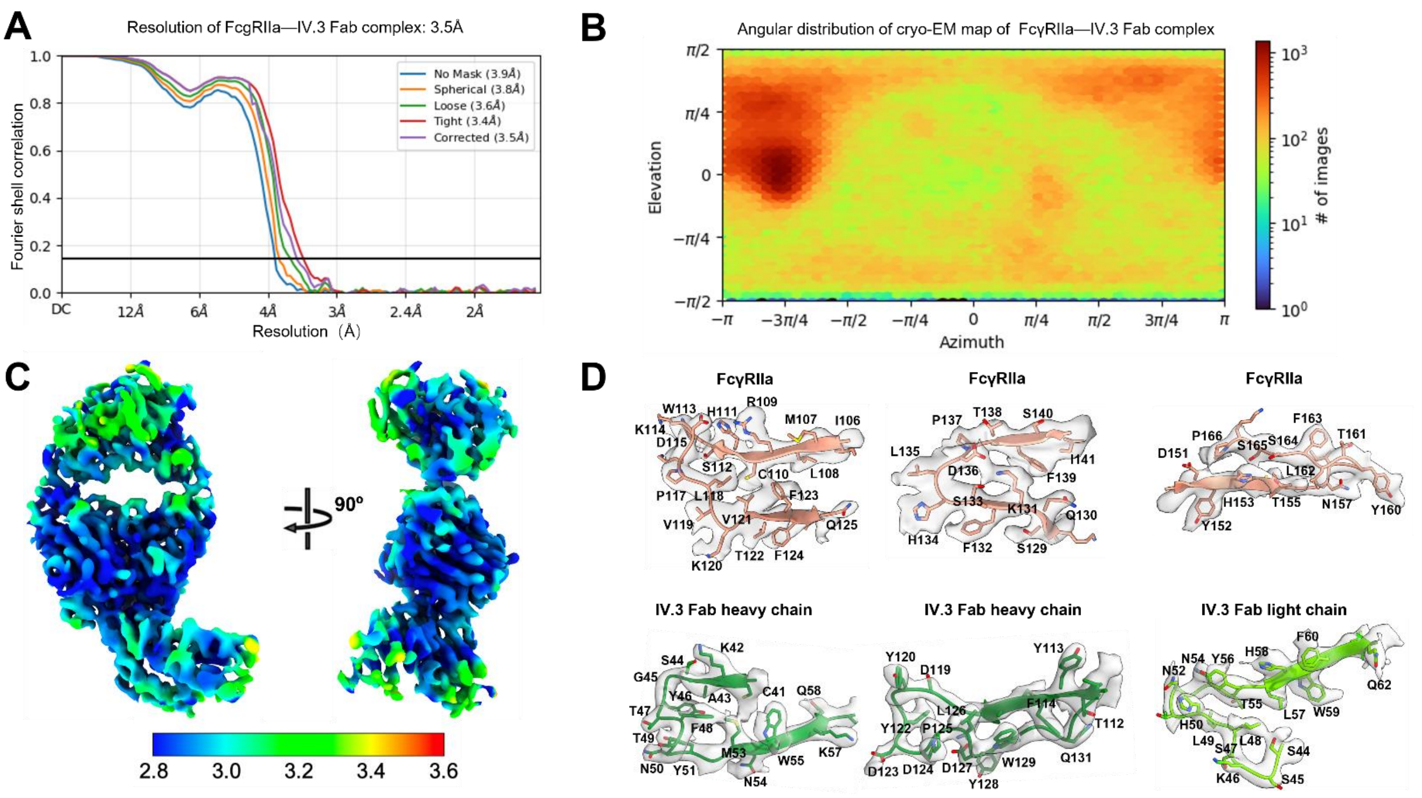


**Supplementary Fig. S3. Cryo–EM reconstruction quality assessment for the FcγRIIa–IV.3 Fab complex.** **(A)** Fourier shell correlation (FSC) curves calculated between independently refined half–maps of the FcγRIIa–IV.3 Fab reconstruction. **(B)** Angular distribution of particle orientations determined in CryoSPARC. **(C)** Local–resolution map of the FcγRIIa–IV.3 Fab reconstruction. **(D)** Representative local cryo–EM density (gray, transparent surface) with the fitted atomic model (colored ribbon representation) showing FcγRIIa and the IV.3 Fab heavy and light chains.

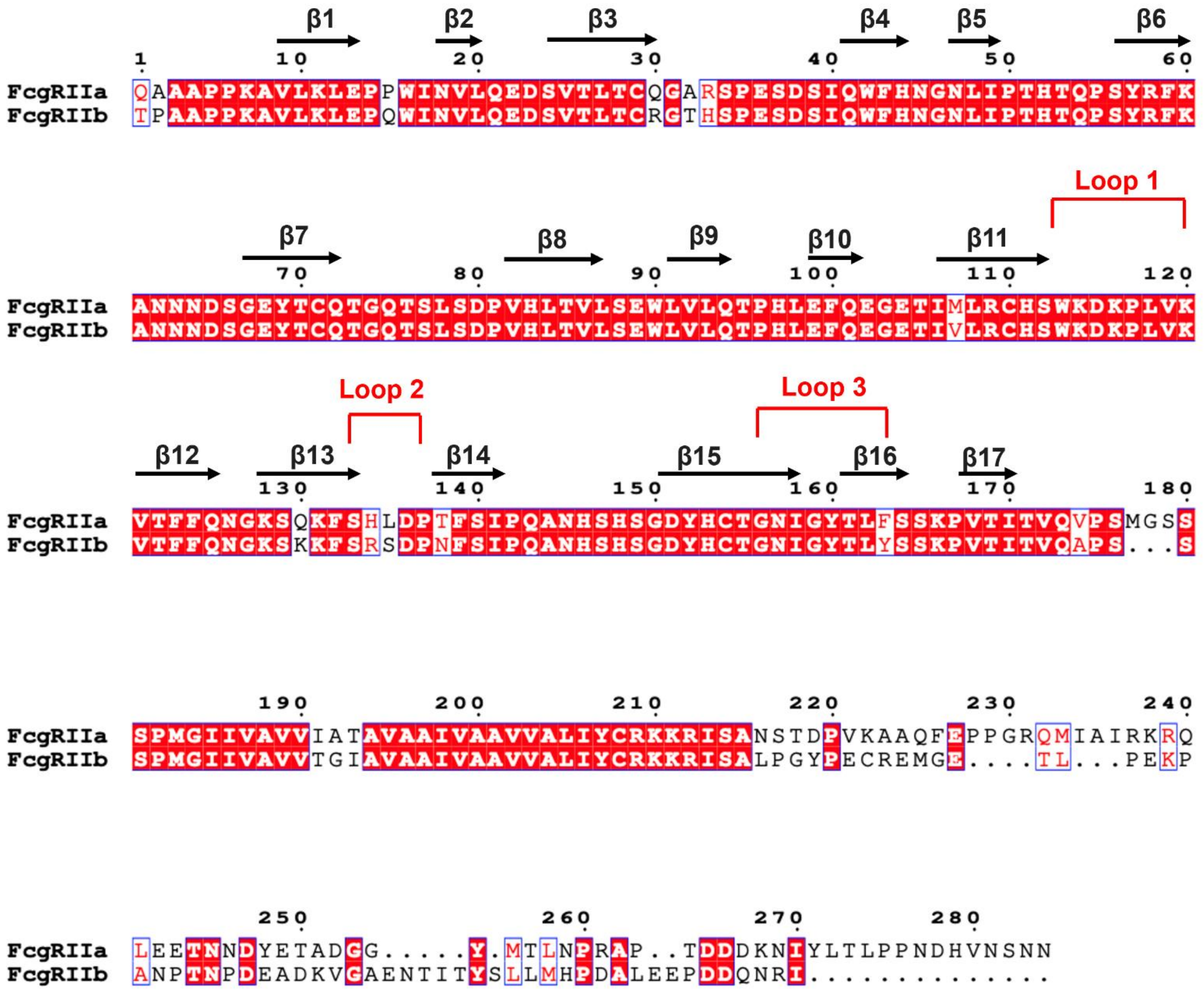

**Supplementary Fig. S4. Sequence comparison between FcγRIIa and FcγRIIb.** Sequence alignment of FcγRIIa and FcγRIIb, with the three loops comprising the IV.3 epitope indicated by red brackets above the corresponding sequence regions. β–strands are annotated and numbered according to their position in the sequence.

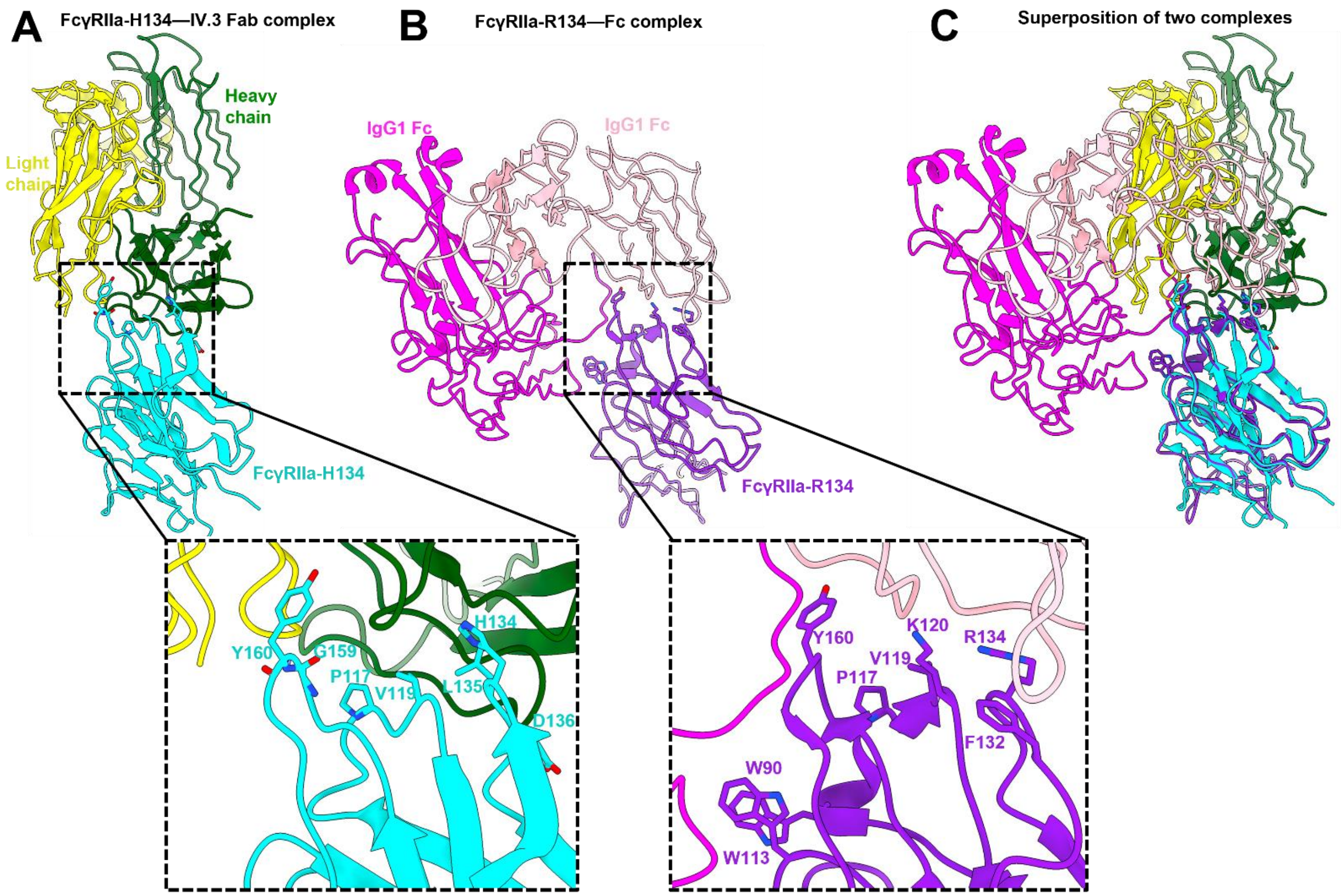


**Supplementary Fig. S5. Comparison of the FcγRIIa–IV.3 Fab cryo–EM structure with the previously published crystal structure of the FcγRIIa–R134 variant bound to the Fc domain of human IgG1. (A)** Cryo–EM structure of the FcγRIIa–H134 variant (cyan) in complex with the IV.3 Fab, with the heavy and light chains shown in green and yellow, respectively. The inset highlights an enlarged view of the interaction interface; FcγRIIa residues identified as interacting by ProLIF analysis (see Supplementary Table 2) are shown as sticks. **(B)** Crystal structure of the FcγRIIa–R134 variant (violet) in complex with the Fc domain of human IgG1 (magenta and pink) (PDB ID: 3RY6). The inset shows an enlarged view of the interaction interface; FcγRIIa residues involved in interactions by ProLIF analysis (see Supplementary Table 2) are shown as sticks. **(C)** Structural superposition of the two complexes following alignment of the FcγRIIa backbones.

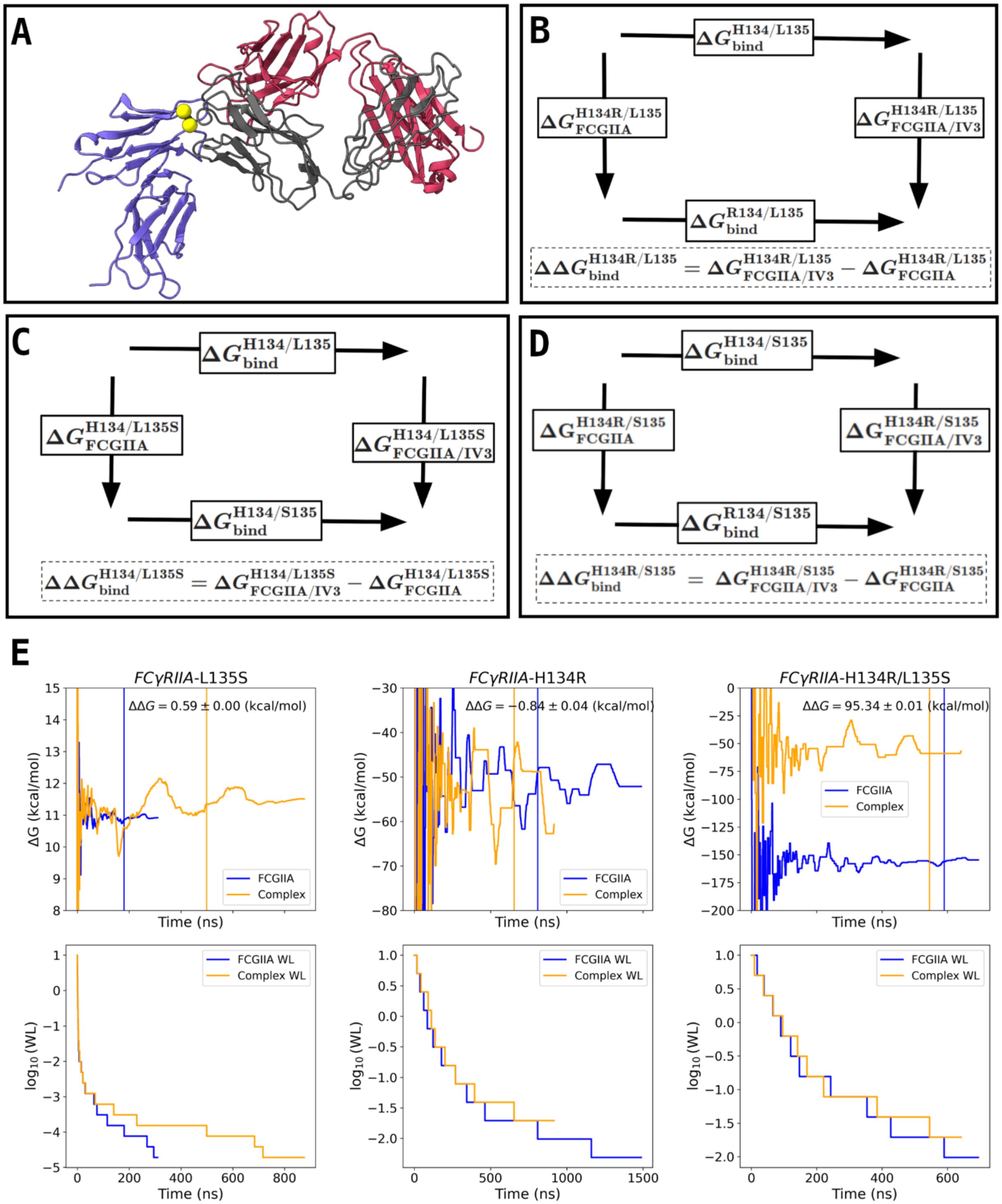


**Supplementary Fig. S6. Theoretical framework for alchemical free energy calculations. (A)** Cryo–EM structure of the FcγRIIa (blue)–IV.3 Fab (red and gray) complex, with variable residues 134 and 135 highlighted as yellow spheres. **(B–D)** Thermodynamic cycles used to compute free–energy changes for the H134R (B), L135S (C), and combined H134R/L135S (D) substitutions. **(E)** Free energy trajectories and Wang–Landau increment trajectories as a function of time.

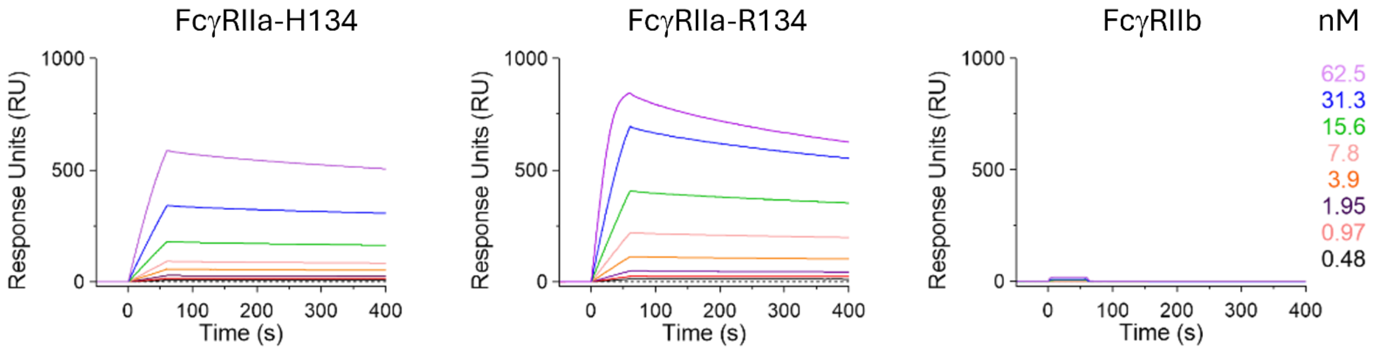


**Supplementary Fig. S7. Surface plasmon resonance analysis of FcγRII binding to mAb IV.3.** Surface plasmon resonance (SPR) sensorgrams of the ectodomains of FcγRIIa-H134, FcγRIIa-R134, and FcγRIIb to immobilized monoclonal antibody IV.3. IV.3 was immobilized on a protein G–coated sensor chip at 10 μg $ml^{-1}$, after which increasing concentrations of each FcγRII variant were injected over the surface. Sensorgrams were globally fit using a 1:1 Langmuir binding model to derive the equilibrium dissociation constants ($K_D$) reported in the main text.

**Supplementary Table S1.** Cryo–EM data collection, refinement and validation statistics.

| | |
|---|---|
| **<u>Data collection</u>** | |
| Microscope | Titan Krios |
| Detector | K3 |
| Voltage (kV) | 300 |
| Electron exposure ($e^{-}/Å^{2}$) | 54 |
| Magnification (kx) | 81 |
| Pixel size (Å) | 0.86 |
| Defocus range (μm) | -1.2 to -2.5 |
| Micrographs collected | 8,898 |
| **Reconstruction** | |
| Symmetry imposed | C1 |
| Initial particle images (No.) | 2,308,718 |
| Final particle images (No.) | 392,060 |
| Box size (px) | 280 |
| Map resolution (Å) | 3.5 |
| FSC threshold | 0.143 |
| Map sharpening B factor ($Å^{2}$) | 218.4 |
| **Model composition** | |
| Non-hydrogen atoms | 0 |
| Protein residues | 612 |
| Metals | 0 |
| **<u>Refinement</u>** | |
| Model-to-map CC (mask) | 0.82 |
| Model-to-map CC (volume) | 0.79 |
| B factor ($Å^{2}$) (min/max/mean) | 9.69/177.91/80.19 |
| Bond r.m.s. deviations | |
| Bond length (Å) | 0.003 |
| Bond angles (º) | 0.612 |
| **<u>Validation</u>** | |
| MolProbity score | 2.52 |
| Clash score | 6.36 |
| Ramachandran plot | |
| Favored (%) | 86.95 |
| Allowed (%) | 12.87 |
| Outliers (%) | 0.17 |
| Rotamer outliers (%) | 5.2 |

**Supplementary Table S2.** FcgRIIa–IV.3 and FcgRIIa–IgG1 (3RY6) protein–protein interaction fingerprints.

| FcγRIIa | IV.3 | IgG1 (3RY6) | Hydrophobic | vdW | Anionic | H-Bond Donor | Face-to-Face | π-Stacking |
|---|---|---|---|---|---|---|---|---|
| W90 | - | G7.A | | X | | | | |
| W90 | - | L98.A | X | X | | | | |
| W90 | - | P99.A | X | | | | | |
| W113 | - | A97.A | | X | | | | |
| W113 | - | P99.A | X | X | | | | |
| P117 | I78.H, Y118.L | L5.A | X | X | | | | |
| P117 | W69.H, Y120.H | - | X | | | | | |
| V119 | D119.H | D35.B, G7.B | | X | | | | |
| V119 | Y120.H | - | X | | | | | |
| K120 | - | D35.B | X | X | X | | | |
| K120 | - | S9.B, V34.B | | X | | | | |
| F132 | - | N67.B | X | X | | | | |
| H134 | D119.H | - | | X | | | | |
| H134 | Y122.H | - | X | X | | | | |
| R134 | - | D35.B | | X | X | | | |
| R134 | - | V36.B | | X | | | | |
| L135 | W69.H | - | X | | | | | |
| L135 | N71.H, Y73.H | - | | X | | | | |
| D136 | Y73.H | - | X | | | | | |
| G159 | Y120.H | - | | X | | | | |
| Y160 | - | L4.B | X | X | | | | |
| Y160 | H50.L | - | X | X | | X | | |
| Y160 | Y56.L | - | X | X | | | X | X |
| Y160 | H115.L | - | X | X | | | | |
| Y160 | Y120.H | - | X | | | | | |

**Supplementary Table S3.** Mutational free–energy estimates of the FcγRIIa–IV.3 Fab complex.

| System | ΔΔG (kcal/mol) | SEM (kcal/mol) |
|---|---|---|
| H134R | -0.84 | 0.04 |
| L135S | 0.59 | 0.00 |
| H134R/L135S | 95.34 | 0.01 |

**Supplementary Table S4.** Net delta contacts per FcγRIIa epitope residue.

| Residue | $H134_{CC1}$ | $H134_{CC2}$ | $H134_{CC3}$ | $R134_{CC1}$ | $R134_{CC2}$ | $R134_{CC3}$ | R134/ $S135_{CC1}$ | R134/ $S135_{CC2}$ | R134/ $S135_{CC3}$ | R134/ $S135_{CC4}$ |
|---|---|---|---|---|---|---|---|---|---|---|
| F132 | 0 | 1 | 0 | 0 | 1 | 0 | 0 | 0 | 0 | 0 |
| S133 | 1 | 1 | 0 | 1 | 0 | 0 | 0 | 0 | 0 | 0 |
| H/R134 | -1 | 4 | 0 | 2 | 3 | 0 | -1 | 0 | 3 | 1 |
| L/S135 | -4 | 0 | 2 | -1 | 2 | -3 | -4 | -6 | -3 | -6 |
| D136 | 1 | 0 | 2 | 1 | 2 | 0 | 2 | -1 | 0 | -1 |
| P137 | 1 | 1 | 2 | 1 | 2 | 1 | 2 | 0 | 2 | 0 |

**Supplementary Table S5.** Initial and final lambdas.

*Initial lambdas for X2R and initial/final lambdas for L2S :*

[0.0 0.1 0.2 0.3 0.4 0.5 0.6 0.7 0.8 0.9 1.0]

*Final lambdas for X2R:*

[0.00000 0.00420 0.00853 0.01300 0.01762 0.02239 0.02734 0.03247 0.03781 0.04337 0.04917 0.05523 0.06158 0.06826 0.07529 0.08271 0.09057 0.09892 0.10780 0.11726 0.12737 0.13814 0.14961 0.16173 0.17442 0.18751 0.20077 0.21400 0.22715 0.24023 0.25323 0.26615 0.27899 0.29175 0.30443 0.31704 0.32962 0.34220 0.35482 0.36752 0.38033 0.39330 0.40648 0.41993 0.43373 0.44799 0.46285 0.47850 0.49523 0.51339 0.53277 0.55242 0.57125 0.58855 0.60411 0.61791 0.63005 0.64075 0.65028 0.65886 0.66665 0.67379 0.68040 0.68654 0.69229 0.69769 0.70280 0.70767 0.71235 0.71689 0.72131 0.72565 0.72992 0.73415 0.73835 0.74254 0.74673 0.75094 0.75519 0.75949 0.76387 0.76834 0.77294 0.77770 0.78267 0.78792 0.79355 0.79972 0.80669 0.81480 0.82462 0.83721 0.85486 0.88052 0.90571 0.92292 0.93527 0.94495 0.95298 0.95988 0.96598 0.97146 0.97645 0.98105 0.98532 0.98931 0.99307 0.99663 1.00000]

**Supplementary Table S6**. Alchemical Expanded Ensemble simulation setup

| System | Box ($nm^3$) | Total Atoms | Total Water | Salt Conc. (M) |
|---|---|---|---|---|
| H134R (complex) | 13.3 | 235,026 | 75,099 | 0.15 |
| H134R (non-complex) | 8.3 | 56,835, | 18,014 | 0.15 |
| L135S (complex) | 13.3 | 235,003 | 75096 | 0.15 |
| L135S (non complex) | 8.3 | 56,848, | 18023 | 0.15 |
| H134R/L135S-(complex) | 13.3 | 235,033 | 75,104 | 0.15 |
| H134R/L135S(non-complex) | 8.3 | 56,830 | 18,015 | 0.15 |

**Supplementary Table S7**. MD simulation setup

| System | Box ($nm^3$) | Total Atoms | Total Water | Salt Conc. (M) |
|---|---|---|---|---|
| H134 | 13.3 | 234,828 | 75,038 | 0.15 |
| R134 | 13.3 | 234,864 | 75,048 | 0.15 |
| R134/S135 | 13.3 | 234,859 | 75,049 | 0.15 |

**Supplementary Movie 1. mAb IV.3 Fab Heavy Chain Loop 1 Rearrangement Around FcγRIIa–R134.** This movie of the molecular dynamics simulation of the FcγRIIa–R134 variant in complex with the mAb IV.3 Fab illustrates a conformational rearrangement of the IV.3 heavy chain Y122 around FcγRIIa R134. In the initial frame of the movie, IV.3 heavy chain residues Y122 and Y51 adopt a gate-like conformation. Over the course of the trajectory, FcγRIIa R134 interacts with Y122 through a cation-π interaction, pulling Y122 away from Y51. R134 then drives Y122 inward towards Y51, distorting the D119-D123 loop. Subsequently, Y122 slides over R134 and flips, relieving loop strain. In the final state, R134 is stabilized in a 'sandwiched' conformation between Y122 and Y51 through cation- π interactions, while also forming a salt-bridge with D119.
